# Evaluation of antiviral drugs against newly emerged SARS-CoV-2 Omicron subvariants

**DOI:** 10.1101/2023.03.26.533897

**Authors:** Junhyung Cho, Younmin Shin, Jeong-Sun Yang, Jun Won Kim, Kyung-Chang Kim, Joo-Yeon Lee

## Abstract

The ongoing emergence of SARS-CoV-2 Omicron subvariants and their rapid worldwide spread pose a threat to public health. From November 2022 to February 2023, newly emerged Omicron subvariants, including BQ.1.1, BF.7, BA.5.2, XBB.1, XBB.1.5, and BN.1.9, became prevalent global strains (>5% global prevalence). These Omicron subvariants are resistant to several therapeutic antibodies. Thus, the antiviral activities of current drugs such as remdesivir, molnupiravir, and nirmatrelvir, which target highly conserved regions of SARS-CoV-2, against newly emerged Omicron subvariants need to be evaluated. We assessed the antiviral efficacy of the drugs using half maximal inhibitory concentration (IC_50_) against human isolated 23 Omicron subvariants and four former SARS-CoV-2 variants of concern (VOC) and compared them with the antiviral efficacy of these drugs against the SARS-CoV-2 reference strain (hCoV/Korea/KCDC03/2020). Maximal IC_50_ fold changes of remdesivir, molnupiravir, and nirmatrelvir were 1.9- (BA.2.75.2), 1.2-(B.1.627.2), and 1.4-fold (BA.2.3), respectively, compared to median IC_50_ values of the reference strain. Moreover, median IC_50_-fold changes of remdesivir, molnupiravir, and nirmatrelvir against the Omicron variants were 0.96, 0.4, and 0.62, similar to 1.02, 0.88, and 0.67, respectively, of median IC_50_-fold changes for previous VOC. Although K90R and P132H in Nsp 5, and P323L, A529V, G671S, V405F, and ins823D in Nsp 12 mutations were identified, these amino acid substitutions did not affect drug antiviral activity. Altogether, these results indicated that the current antivirals retain antiviral efficacy against newly emerged Omicron subvariants, and provide comprehensive information on the antiviral efficacy of these drugs.

## Text

The continuous emergence of SARS-CoV-2 Omicron subvariants is accompanied by rapid spread worldwide. Since its appearance in November 2021, the Omicron variant, B.1.1.529, has outcompeted other variants of concern (VOC) with 26–32 amino acid changes in the spike protein, and Omicron subvariants have become the dominant strains worldwide (Guo et al., 2022). The globally prevalent Omicron sublineages (>5% global preference) from November 2022 to February 2023 were BQ.1.1, BF.7, BA.5.2, XBB.1, XBB.1.5, and BN.1.9 variants (Khare et al., 2021). The emergence of these Omicron subvariants is concerning because of the possibility of resistance to current antivirals.

Mainly, two classes of targeted antiviral drugs have been developed for treating COVID-19. The first are monoclonal antibodies, which bind directly to the SARS-CoV-2 spike protein and inhibits viral infection (Brady et al., 2022). Although several monoclonal antibodies showed effective virus neutralisation early in the pandemic, they were less or not effective against the emerging Omicron subvariants (Arora et al., 2023).

The second are small-molecule drugs that target viral RNA-dependent RNA polymerase (RdRp) or main protease (Mpro) (Brady et al., 2022). Remdesivir (Veklury) and molnupiravir (Lagevrio) are nucleoside analogue prodrugs targeting RdRp of SARS-CoV-2, and nirmatrelvir/ritonavir (Paxlovid) is a peptidomimetic inhibitor of the SARS-CoV-2 Mpro. Remdesivir became the first US Food and Drug Administration (FDA) approved drug in October 2020, and molnupiravir and Nirmatrelvir/ritonavir received emergency use authorisation by the FDA in December 2021 (Brady et al., 2022). Despite these drugs targeting highly conserved regions of the virus genome, genetic mutations could occur in the spike protein and elsewhere in the viral genome (Kim et al., 2020). Thus, the antiviral efficacy of the drugs used for COVID-19 patients need to be assessed against overall SARS-CoV-2 variants.

In this study, we assessed the antiviral efficacy of the drugs by comparing the foldchange ratio of the *in vitro* half maximal inhibitory concentration (IC_50_) and median IC_50_ value against human isolated 23 new Omicron subvariants and four former SARS-CoV-2 VOCs to that of the reference strain (hCoV/Korea/KCDC03/2020), which showed high homology of >99.5% with first isolated SARS-CoV-2 sequence in Wuhan, China (Kim et al., 2020).

To measure the IC_50_ of the drugs against authentic virus infection, 0.1 multiplicity of infection of each virus was inoculated onto Vero E6 cells (ATCC, Manassas, VA, USA). Subsequently, seven concentrations of the drugs were treated with two-fold serial dilution [remdesivir and nirmatrelvir (antiviral component of Paxlovid): 20 to 0.31 μM, molnupiravir: 40 to 0.62 μM] for 48 h. The cell infectivity ratio between the drug treatment and virus only-treated groups was assessed through high-content imaging (HCI) analysis to determine the number of infected cells (N protein expressed cells) using immunofluorescence images with viral N-specific antibody and total cells (number of nuclei) using DAPI staining (Supplementary Figure S1A; the methods described in the Supplementary Information.) The IC_50_ and 95% confidence interval (CI) values were determined from dose-response curves based on treatment with seven concentrations of each drug, using Prism 7 (GraphPad Software, San Diego, CA, USA), and the quality of the assay was assessed using a Z’ factor >0.5 (Table 1 and Supplementary Figure S1BC).

**Table 1.**
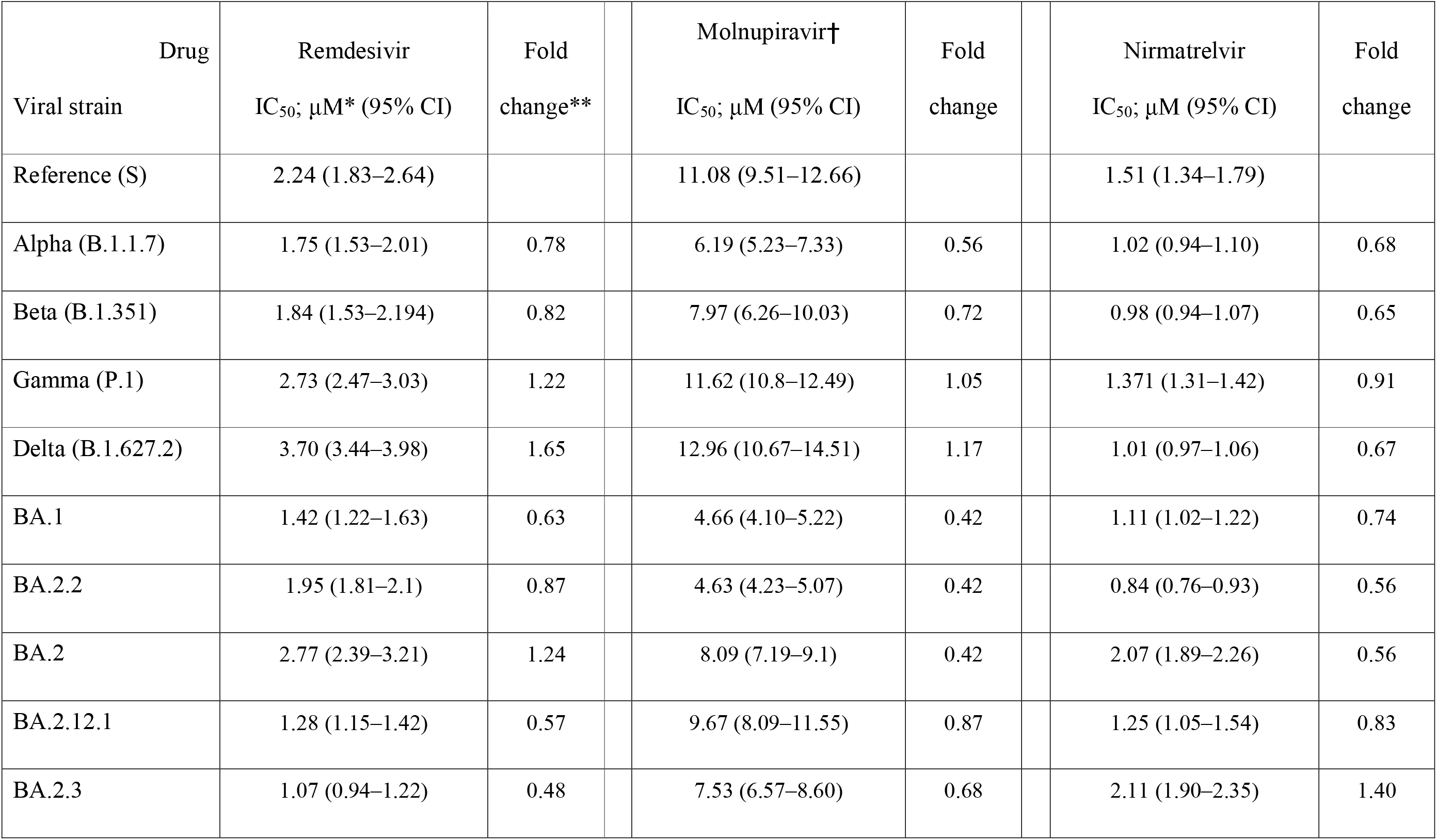
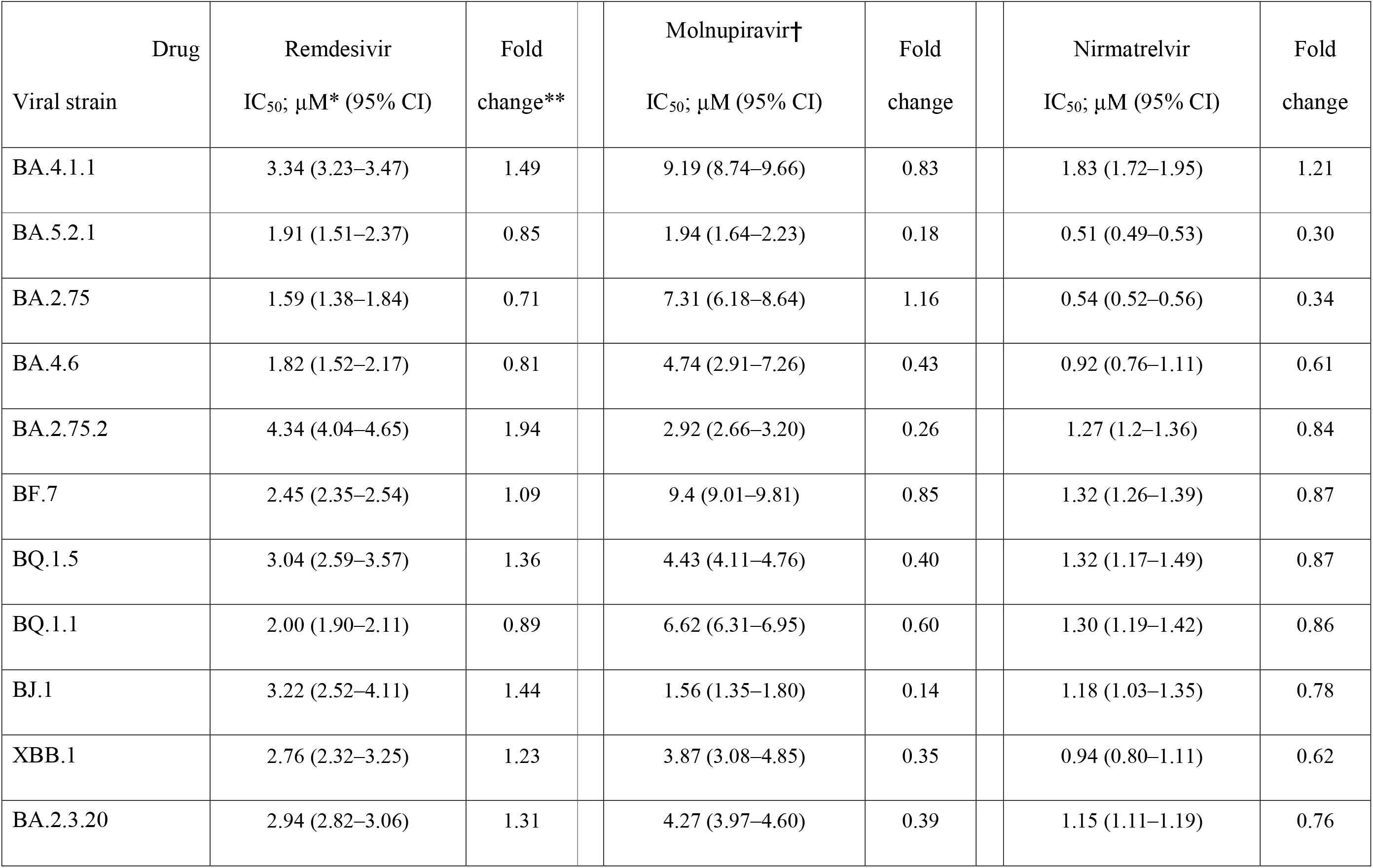
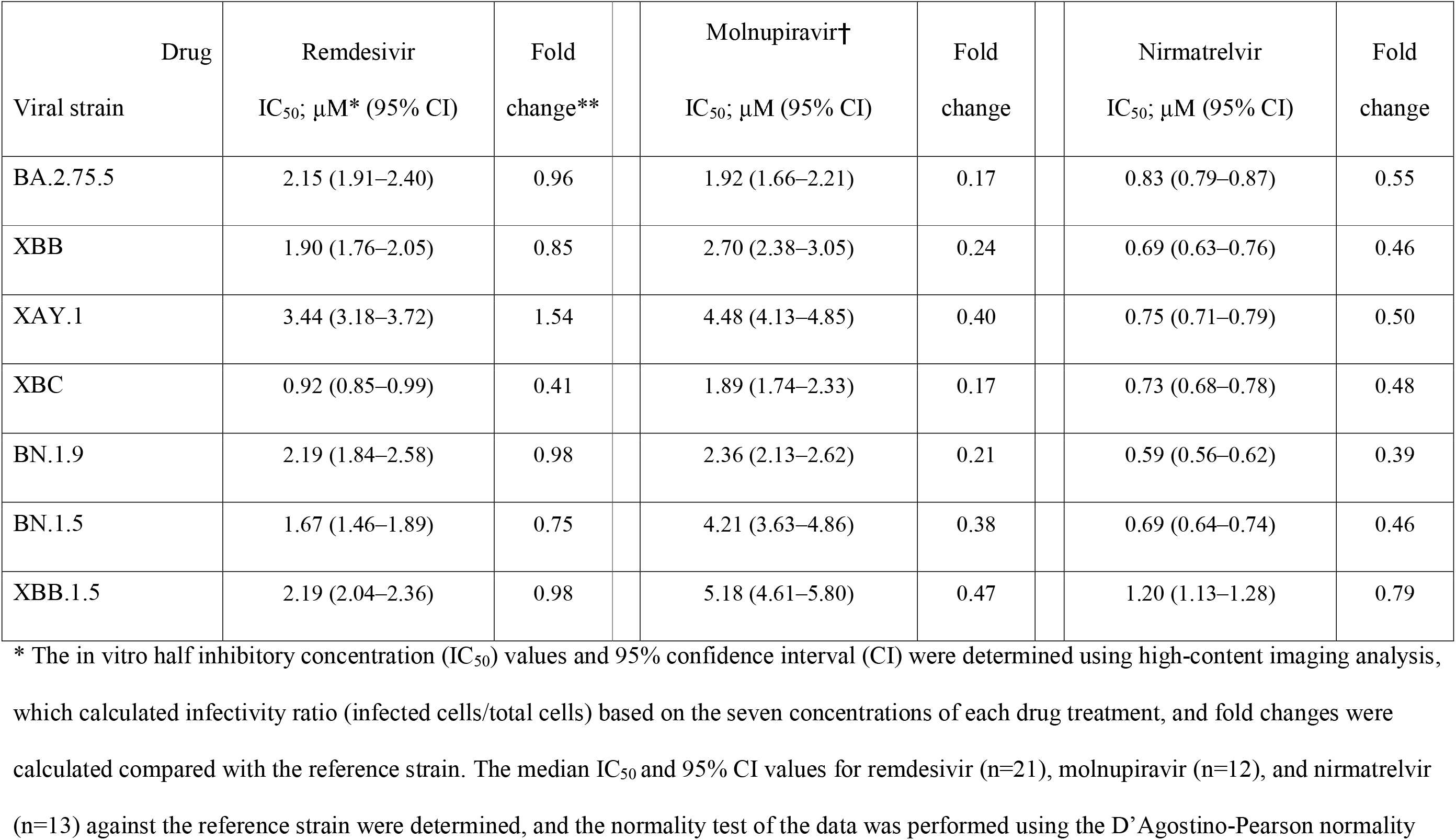
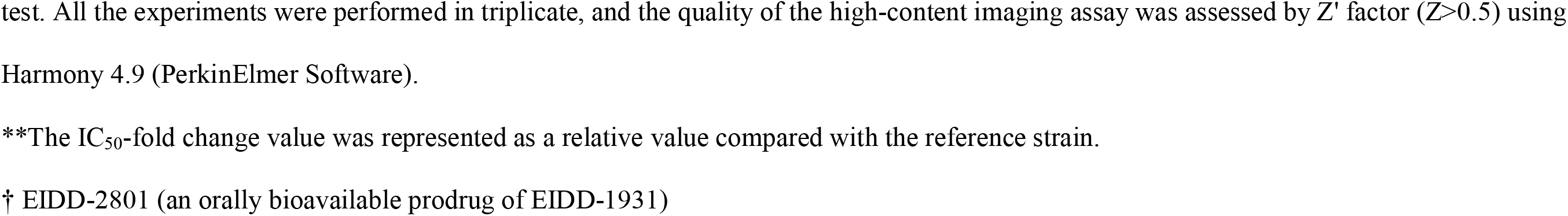
IC_50_ values of remdesivir, molnupiravir, and nirmatrelvir against SARS-CoV-2 variants.

The maximal IC_50_-fold change values of remdesivir, molnupiravir, and nirmatrelvir against all the SARS-CoV-2 variants were 1.9-(BA.2.75.2), 1.2-(B.1.627.2), and 1.4-fold (BA.2.3), respectively, compared to the median IC_50_ values of the reference strain for remdesivir, molnupiravir, and nirmatrelvir (Figure 1 and Table 1). Moreover, the median IC_50_-fold change values in remdesivir, molnupiravir, and nirmatrelvir for the Omicron variants were 0.96-, 0.4-, and 0.62-fold, respectively, which were similar to the median IC_50_-fold change values of 1.02, 0.88, and 0.67, respectively, for the former VOC (Table 1).

**Figure 1.**
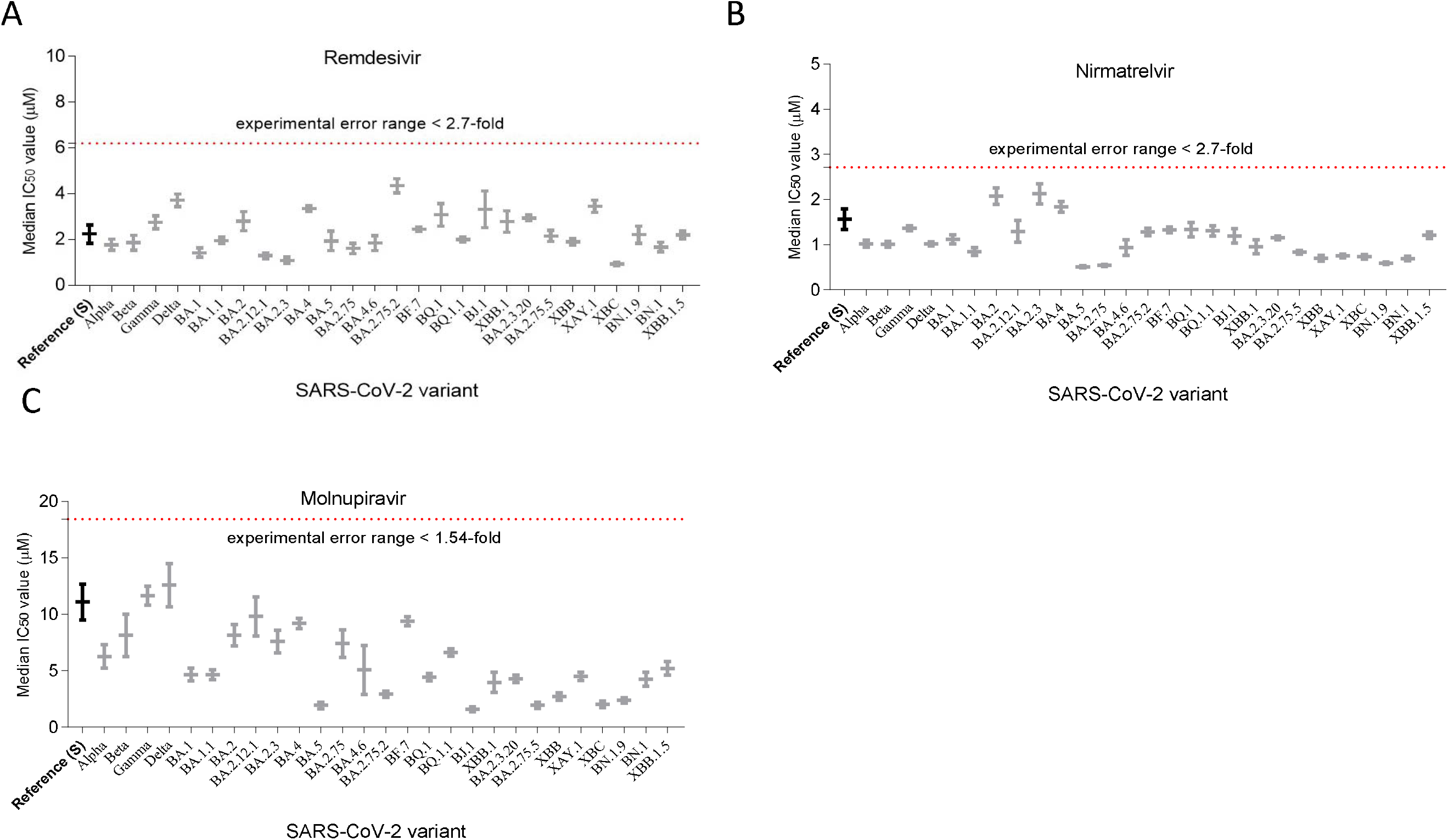
Median IC50 value of the drugs in VeroE6 cells. Median IC50 (left axis) of (A) remdesivir, (B) nirmatrelvir, and (C) molnupiravir. Vero E6 cells were infected with indicated SARS-CoV-2 variants at 0.1 multiplicity of infection for 1 h. Subsequently, seven concentrations of the drugs were treated with two-fold serial dilution [remdesivir and nirmatrelvir (antiviral component of Paxlovid): 20 to 0.31 μM, molnupiravir: 40 to 0.62 μM] for 48 h. The cell infectivity ratio between the drug treatment and virus only-treated groups was assessed through high-content imaging analysis to determine the number of infected cells (N protein expressed cells) using immunofluorescence images with viral N-specific antibody and total cells (number of nuclei) using DAPI staining. The IC_50_ values were determined from dose-response curves and median IC_50_ value was calculated. The experiments were performed in triplicate [the reference strain for remdesivir (n=21), nirmatrelvir (n=13), and molnupiravir (n=12)].

Using GISAID sequence analysis, BA.2.75.2 and Delta (B.1.627.2) have P323L and G671S amino acid substitution in nonstructural protein (Nsp) 12, and BA.2.3 has P132H amino acid substation in Nsp 5 (Supplementary Table S1). However, it was reported that high prevalent changes, P323L (>99% in Omicron variant) and G671S (>99% in Delta variant) substitution in Nsp 12 and P132H substitution in Nsp 5 did not affect the antiviral activity of nirmatrelvir due to the distance of drug binding location (Pitts et al., 2022; Ullrich et al., 2022). Moreover, the maximal fold changes of the drugs were within experimental error range in our experiment condition in which the 95% and 5% percentile ratio of the IC_50_ for remdesivir (n=21), molnupiravir (n=13), and nirmatrelvir (n=12) on the reference strain were approximately 2.7-, 1.5-, and 1.9-fold, respectively (Supplementary Figure S1B) (Vangeel et al., 2022).

Other amino acid substitutions such as V405F (BA.2.75), Y273H (BQ.1.1 and BQ.1.5), A529V (XAY.1 and XBC), and ins823D (XBC) were identified in Nsp 12. A529V, V405F, X823D, and Y273H substitutions were previously reported, where their frequency was approximately 1%, and did not affect the binding of remdesivir (Pitts et al., 2022; Vangeel et al., 2022; Kumar et al., 2022; Imai et al., 2023). Moreover, they did not include the most important residues for RdRp binding (Nguyen et al., 2020). In Nsp 5, K90R substitutions were previously reported in Beta (B.1.351) and XBB; however, the substitution did not change the antiviral activity of nirmatrelvir (Imai et al., 2023; Hu et al., 2022). Thus, the drugs could retain their antiviral activity against newly emerging Omicron subvariants.

In our results, IC_50_ value of the drugs was approximately 10-fold higher than that in other similar studies (Ullrich et al., 2022; Vangeel et al., 2022). In drug treatment, EIDD-2801 (an orally bioavailable prodrug of EIDD-1931) and EIDD-1931 (an active form of molnupiravir) treatment in Vero E6 cells did not show significant IC_50_ value changes (Vangeel et al., 2022). Thus, it may be due to without treatment of P-glycoprotein inhibitor (P-pg, drug efflux inhibitor), which enhanced IC_50_ values by 10–100-fold for nirmatrelvir and remdesivir in Vero E6 cells and experimental conditions (Vangeel et al., 2022; Imai et al., 2023; Fiaschi et al., 2022; Zhu et al., 2022).

Altogether, we assessed the antiviral activity of these drugs against newly emerging Omicron subvariants and confirmed that it was maintained. Moreover, the mutations of Nsp 5 and Nsp 12 in the Omicron subvariants did not affect the antiviral activity of the drugs. Our results provide comprehensive information that antiviral efficacy of remdesivir, molnupiravir, and nirmatrelvir against 27 SARS-CoV-2 variants is maintained and thus could be used continuously for clinical treatment with COVID-19 patients.

## ^1^Abbreviations

IC_50_: half maximal inhibitory concentration
VOC: variants of concern
FDA: Food and Drug Administration
RdRp: RNA-dependent RNA polymerase
Mpro: main protease
CI: confidence interval

## References

Arora, P., Kempf, A., Nehlmeier, I., Schulz, SR., Jäck, HM., Pöhlmann, S., Hoffmann, M., 2023. Omicron sublineage BQ.1.1 resistance to monoclonal antibodies. Lancet Infect. Dis. 23, 22–23. https://doi.org/10.1016/S1473-3099(22)00733-2.

Brady, D.K., Gurijala, AR., Huang, L., Hussain, AA., Lingan, AL., Pembridge, OG., Ratangee, BA., Sealy, TT., Vallone, KT., Clements, TP., 2022. A guide to COVID□19 antiviral therapeutics: a summary and perspective of the antiviral weapons against SARS□CoV□2 infection. FEBS Journal, 1–31. https://doi.org/10.1111/febs.16662.

Fiaschi, L., Dragoni, F., Schiaroli, E., Bergna, A., Rossetti, B., Giammarino, F., Biba, C., Gidari, A., Lai, A., Nencioni, C., Francisci, D., Zazzi, M., Vicenti, I., 2022. Efficacy of licensed monoclonal antibodies and antiviral agents against the SARS-CoV-2 omicron sublineages BA.1 and BA. Viruses. 14. https://doi.org/10.3390/v14071374.

Guo, Y., Han, J., Zhang, Y., He, J., Yu, W., Zhang, X., Wu, J., Zhang, S., Kong, Y., Guo, Y., Lin, Y., Zhang, J., 2022. SARS-CoV-2 Omicron variant: epidemiological features, biological characteristics, and clinical significance. Front. Immunol. 13, 877101. https://doi.org/10.3389/fimmu.2022.877101.

Hu, Y., Lewandowski, EM., Tan, H., Zhang, X., Morgan, RT., Zhang, X., Jacobs, LMC., Butler, SG., Gongora, MV., Choy, J., Deng, X., Chen, Y., Wang, J., 2022. Naturally occurring mutations of SARS-CoV-2 main protease confer drug resistance to nirmatrelvir. bioRxiv. https://doi.org/10.1101/2022.06.28.497978.

Imai, M., Ito, M., Kiso, M., Yamayoshi, S., Uraki, R., Fukushi, S., Watanabe, S., Suzuki, T., Maeda, K., Sakai-Tagawa, Y., Iwatsuki-Horimoto, K., Halfmann, PJ., Kawaoka, Y., 2023. Efficacy of antiviral agents against omicron subvariants BQ.1.1 and XBB. N. Engl. J. Med. 388, 89–91. https://doi.org/10.1056/NEJMc2214302.

Khare, S., Gurry, C., Freitas, L., Schultz, MB., Bach, G., Diallo, A., Akite, N., Ho, J., Lee, RT., Yeo, W., Curation Team GC., Maurer-Stroh S., 2021. GISAID’s role in pandemic response. China CDC Wkly. 3, 1049–1051. https://doi.org/10.46234/ccdcw2021.255.

Kim, J.M., Jang, JH., Kim, JM., Chung, YS., Yoo, CK., Han, MG., 2020. Identification of coronavirus isolated from a patient in Korea with COVID-19. Osong Public Health Res. Perspect. 11, 3–7. https://doi.org/10.24171/j.phrp.2020.11.1.02.

Kim, J.S., Jang, JH., Kim, JM., Chung, YS., Yoo, CK., Han, MG., 2020. Genome-wide identification and characterization of point mutations in the SARS-CoV-2 genome.Osong Public Health Res. Perspect. 11, 101–111.https://doi.org/10.24171/j.phrp.2020.11.3.05.

Kumar, S., Kumari, K., Azad, GK., 2022. Emerging genetic diversity of SARS-CoV-2 RNA dependent RNA polymerase (RdRp) alters its B-cell epitopes. Biologicals. 75, 29–36. https://doi.org/10.1016/j.biologicals.2021.11.002.

Nguyen, H.L., Thai, NQ., Truong, DT., Li, MS., 2020. Remdesivir strongly binds to both RNA-dependent RNA polymerase and main protease of SARS-CoV-2: evidence from molecular simulations. J. Phys. Chem. B. 124, 11337–11348. https://doi.org/10.1021/acs.jpcb.0c07312.

Pitts, J. Li, J., Perry, JK., Du Pont, V., Riola, N., Rodriguez, L., Lu, X., Kurhade, C., Xie, X., Camus, G., Manhas, S., Martin, R., Shi, PY., Cihlar, T., Porter, DP., Mo, H., Maiorova, E., Bilello, JP., 2022. Remdesivir and GS-441524 retain antiviral activity against Delta, Omicron, and other emergent SARS-CoV-2 variants. Antimicrob.Agents Chemother. 66, e0022222.https://doi.org/10.1128/aac.00222-22.

Ullrich, S., Ekanayake, KB., Otting, G., Nitsche, C., 2022. Main protease mutants of SARS-CoV-2 variants remain susceptible to nirmatrelvir. Bioorg. Med. Chem. Lett. 62, 128629. https://doi.org/10.1016/j.bmcl.2022.128629.

Vangeel, L., Chiu, W., De Jonghe, S., Maes, P., Slechten, B., Raymenants, J., André, E., Leyssen, P., Neyts, J., Jochmans, D., 2022. Remdesivir, molnupiravir and nirmatrelvir remain active against SARS-CoV-2 Omicron and other variants of concern. Antiviral Res. 198, 105252.https://doi.org/10.1016/j.antiviral.2022.105252.

Zhu, Y., Binder, J., Yurgelonis, I., Rai, DK., Lazarro, S., Costales, C., Kobylarz, K., McMonagle, P., Steppan, CM., Aschenbrenner, L., Anderson, AS., Cardin, RD., 2022. Generation of a VeroE6 Pgp gene knock out cell line and its use in SARS-CoV-2 antiviral study. Antiviral Res. 208, 105429.https://doi.org/10.1016/j.antiviral.2022.105429.

