## Supplementary Information for "Evaluation of antiviral drugs against newly emerged SARS-CoV-2 Omicron subvariants"

187, Osongsaengmyeong2-ro, Osong-eup, Heungdeok-gu, Cheongju-si, Chungcheongbuk-do 28159, Republic of Korea

 (KCK); (JYL)

**Abstract**

The ongoing emergence of SARS-CoV-2 Omicron subvariants and their rapid worldwide spread pose a threat to public health. From November 2022 to February 2023, newly emerged Omicron subvariants, including BQ.1.1, BF.7, BA.5.2, XBB.1, XBB.1.5, and BN.1.9, became prevalent global strains (>5% global prevalence). These Omicron subvariants are resistant to several therapeutic antibodies. Thus, the antiviral activities of current drugs such as remdesivir, molnupiravir, and nirmatrelvir, which target highly conserved regions of SARS-CoV-2, against newly emerged Omicron subvariants need to be evaluated. We assessed the antiviral efficacy of the drugs using half maximal inhibitory concentration (IC_50_) against human isolated 23 Omicron subvariants and four former SARS-CoV-2 variants of concern (VOC) and compared them with the antiviral efficacy of these drugs against the SARS-CoV-2 reference strain (hCoV/Korea/KCDC03/2020). Maximal IC_50_ fold changes of remdesivir, molnupiravir, and nirmatrelvir were 1.9- (BA.2.75.2), 1.2- (B.1.627.2), and 1.4-fold (BA.2.3), respectively, compared to median IC_50_ values of the reference strain. Moreover, median IC_50_-fold changes of remdesivir, molnupiravir, and nirmatrelvir against the Omicron variants were 0.96, 0.4, and 0.62, similar to 1.02, 0.88, and 0.67, respectively, of median IC_50_-fold changes for previous VOC. Although K90R and P132H in Nsp 5, and P323L, A529V, G671S, V405F, and ins823D in Nsp 12 mutations were identified, these amino acid substitutions did not affect drug antiviral activity. Altogether, these results indicated that the current antivirals retain antiviral efficacy against newly emerged Omicron subvariants, and provide comprehensive information on the antiviral efficacy of these drugs.

Keywords: SARS-CoV-2, Omicron subvariant, remdesivir, molnupiravir, nirmatrelvir, antiviral activity

**Materials and methods**

**Cell culture and viruses**

SARS-CoV-2 variants and virus information were obtained from the National Culture Collection for Pathogens of the Korea National Institute of Health (Cheongju-si, South Korea). Vero E6 cells were purchased from the American Type Culture Collection (ATCC CCL-81, Manassas, VA, USA) and cultured in Dulbecco’s minimal essential medium (DMEM; Gibco, Grand Island, NY, USA) supplemented with 2% (infection media) or 10% (growth media) v/v heat-inactivated foetal bovine serum (FBS; Gibco) and 1% v/v penicillin/streptomycin (p/s; Gibco) in a humidified incubator with 5% CO_2_ at 37 °C. Remdesivir, molnupiravir (EIDD-2801), and nirmatrelvir were purchased from MedChemExpress (Monmouth Junction, NJ, USA).

**IC_50_ measurement**

Vero E6 cells (1×10^4^ cells/well) were seeded in a cell carrier ultra-96 plate (PerkinElmer, Waltham, MA, USA) 24 h before infection. The next day, 0.1 multiplicity of infection of the virus in 100 µL of infection media (DMEM containing 2% FBS and 1% p/s) was added to cells for 1 h to allow infection, and the cells were then washed with phosphate-buffered saline (PBS; Gibco). Subsequently, seven concentrations of remdesivir and nirmatrelvir (20, 10, 5, 2.5, 1.25, 0.625, and 0.312 μM) and molnupiravir (40, 20, 10, 5, 2.5, 1.25, and 0.625 μM) were added to the cells and incubated for 48 h. After incubation, the supernatant was removed, and the cells were washed twice with PBS and fixed with 4% formaldehyde for 15 min. Cells were washed and incubated for 2 h with anti-SARS-CoV-2 N antibody (1:2,000, Rabbit IgG) (Sinobio Cat#40143-V08B, Beijing, China) in western blocking solution (Sigma-Aldrich, Burlington, MA, USA) and then with Alexa488-conjugated rabbit antibody (1:2,000) (Thermo Fisher Scientific, Waltham, MA, USA) for 1 h. Subsequently, the cells were incubated with DAPI (1:1,000) (Abcam, Waltham, MA, USA) in PBS for 10 min and washed five times with PBS containing Tween (Sigma-Aldrich). Immunofluorescence images were acquired using a 10× objective with five imaged fields per well using Operetta CLS (PerkinElmer). The acquired images were analysed using the Harmony software (ver. 4.9, PerkinElmer) to quantify total cell numbers (nucleus, DAPI) and infected cell numbers (virus N protein expression cells, Alexa488), and cell infectivity was normalised to the virus alone (0.5% DMSO) group and noninfected cells (0.5% DMSO). The cell infectivity ratio (Y-axis) was calculated as follows: viral N protein cell numbers (N protein, Alexa488 expressed cells)/total cell numbers (DAPI stained cells) × 100. Subsequently, the drug concentration (X-axis) was transformed to logarithms with base 10, and nonlinear regression of the log[inhibitor] vs. normalised response IC_50_ equation (Y=100/(1+10^((LogIC_50_−X)*HillSlope))) was employed using Prism 7 (GraphPad software, San Diego, CA, USA) to determine the IC_50_ and 95% confidence interval (CI) of the drugs, where X is the log of drug concentration, Y is normalised response (100% down to 0%), and HillSlope is the slope factor or Hill slope. The IC_50_ and 95% CI values were determined using Prism 7, and the quality of the HCI assay was assessed using the Z' factor (Z' > 0.5) with Harmony 4.9 (PerkinElmer Software). All experiments were performed in triplicate, and authentic virus infection was performed in biosafety level 3 facilities as per the Korea Disease Control and Prevention Agency guidelines.

**Supplementary Figure legend**

Figure S1. Dose-response curve analysis of SARS-CoV-2 variant infections exposed to different concentrations of antiviral drugs. (A) Dose-response curve analysis images of the drug treatment group by immunofluorescence against SARS-CoV-2 reference strain (hCoV/Korea/KCDC03/2020) infection. The microscope images show SARS-CoV-2 N protein (green) and cell nuclei (blue) at the specified drug concentration (scale bar is 1mm). (B) Median IC_50_ values of the drugs for the reference strain infection. The median IC_50_ and 95% confidence interval (CI) values for remdesivir (n=21), molnupiravir (n=12), and nirmatrelvir (n=13) against the reference strain were determined using Prism 7, and the normality test of the data was performed using the D’Agostino-Pearson normality test. (C) Dose-response curve analysis of the drugs. Seven concentrations of each drug were used for treatment, and the cell infectivity of the SARS-CoV-2 variants was calculated using the ratio of N protein positive cell to the total cell numbers. The IC_50_ values were normalised to virus only (DMSO) and non-infection groups, and the IC_50_ values and 95% CIs were determined using Prism 7. All the experiments were performed in triplicate, and the quality of the HCI assay was assessed using the Z' factor (Z' > 0.5) with Harmony 4.9 (PerkinElmer Software).

Supplementary Figure S1

(A)

**Remdesivir**

**0.15**

**0.31**

**0.62**

**1.25**

**2.5**

**5**

**20**

**10**

**Nirmatrelvir**

**Molnupiravir**

**0**

**0**

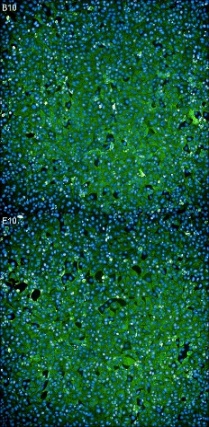

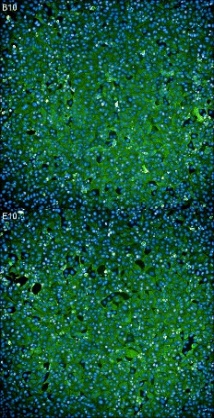

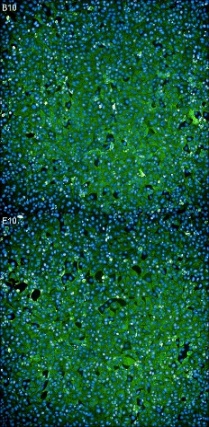

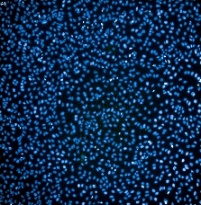

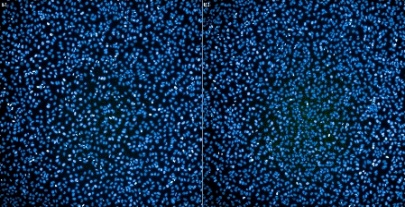

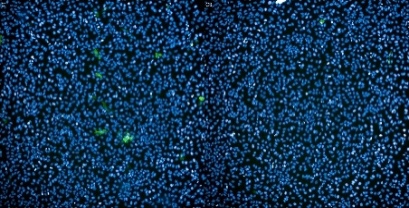

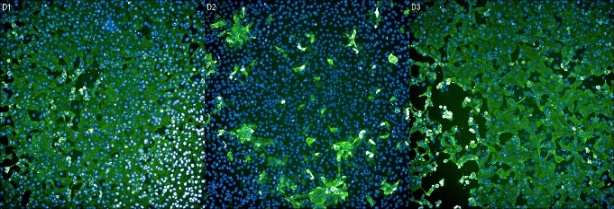

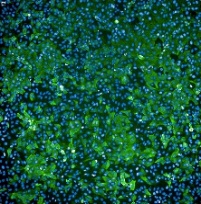

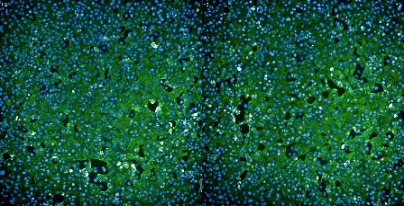

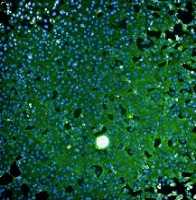

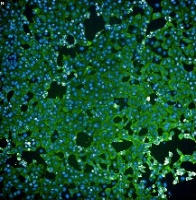

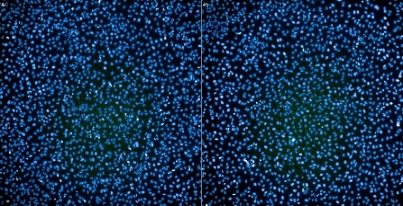

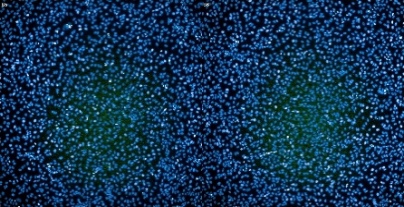

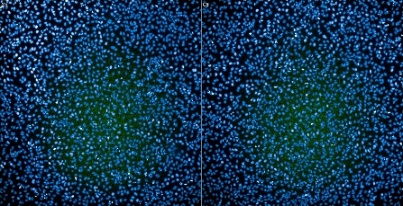

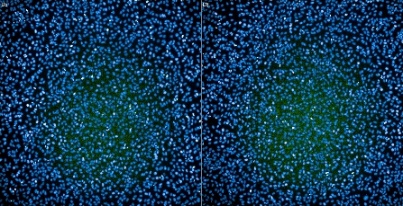

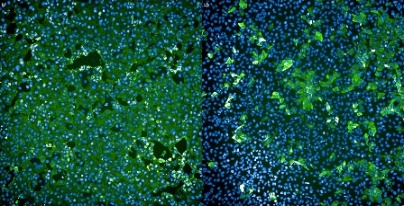

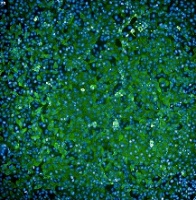

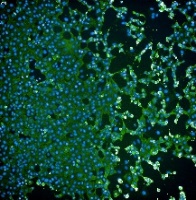

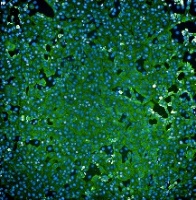

**0**

**0.15**

**0.31**

**0.62**

**1.25**

**2.5**

**5**

**20**

**10**

**0.31**

**0.62**

**1.25**

**2.5**

**5**

**20**

**10**

**40**

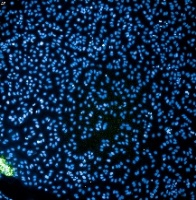

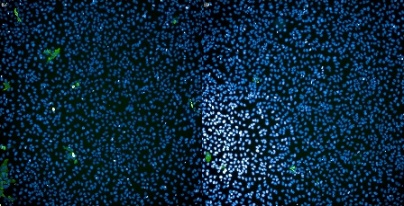

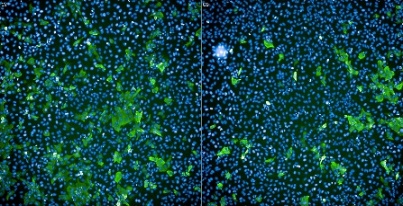

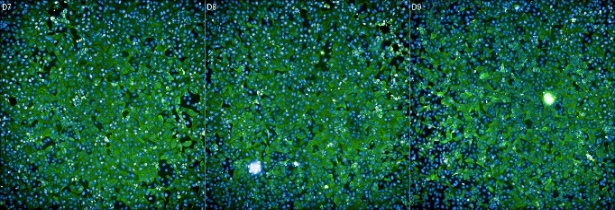

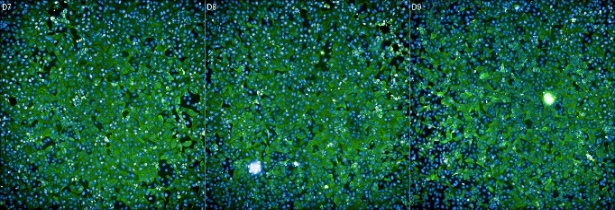

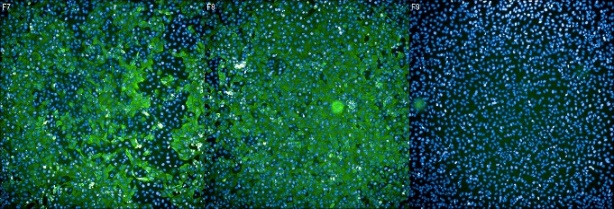

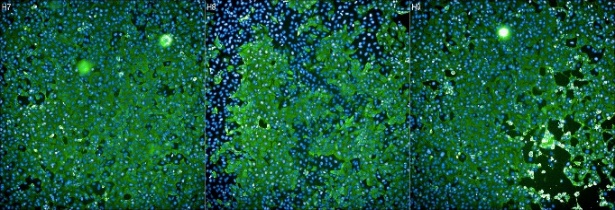

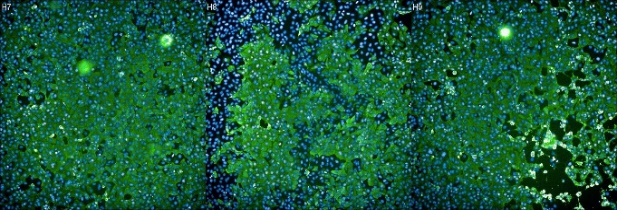

**μΜ:**

**μΜ:**

**μΜ:**

1mm

1mm

1mm

(B)

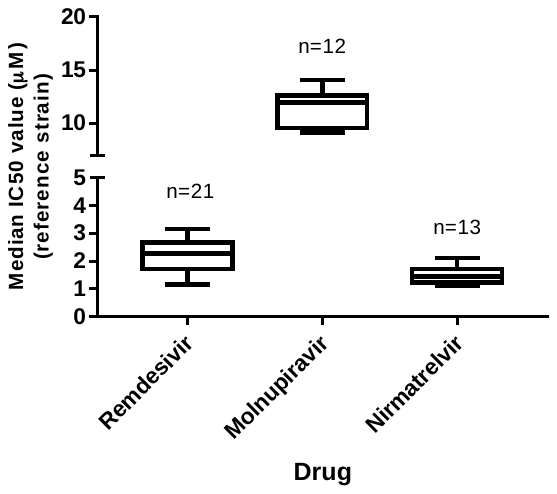

(C)

IC_50_=1.84 μM, R^2^=0.93

IC_50_=0.98 μM, R^2^=0.96

IC_50_=1.75 μM, R^2^=0.96

**Remdesivir**

IC_50_=1.02 μM, R^2^=0.98

**Nirmatrelvir**

**Alpha**

**Beta**

**Molnupiravir**

**Gamma**

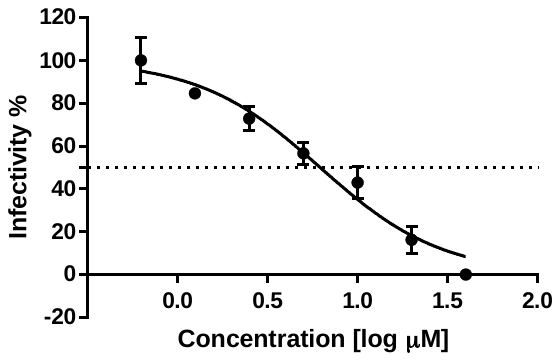

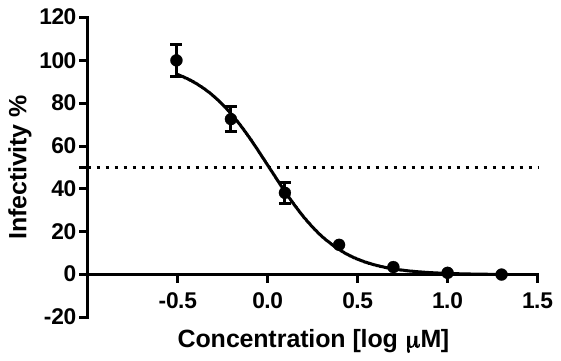

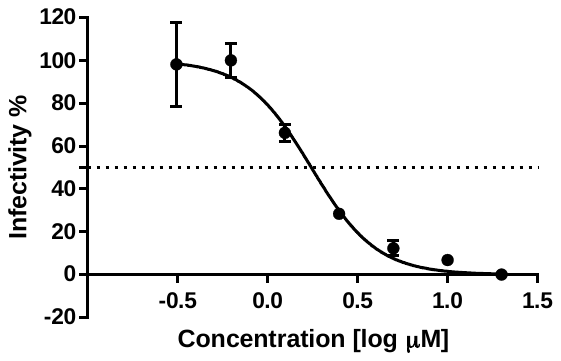

IC_50_=6.19 μM, R^2^=0.95

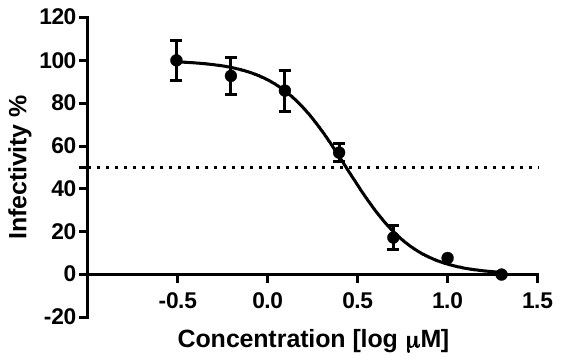

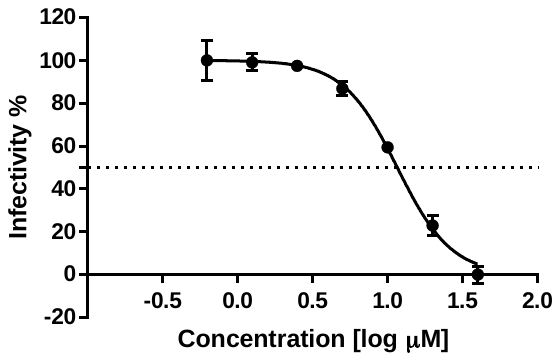

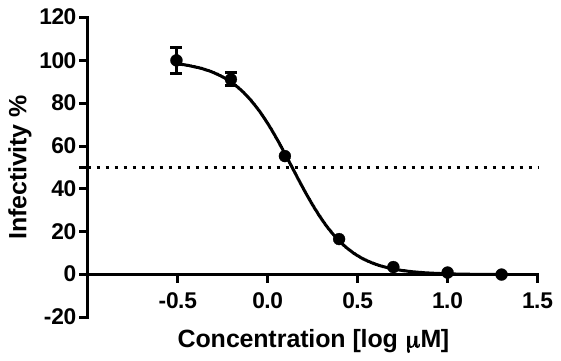

IC_50_=2.73 μM, R^2^=0.97

IC_50_=1.37 μM, R^2^=0.99

IC_50_=11.62 μM, R^2^=0.98

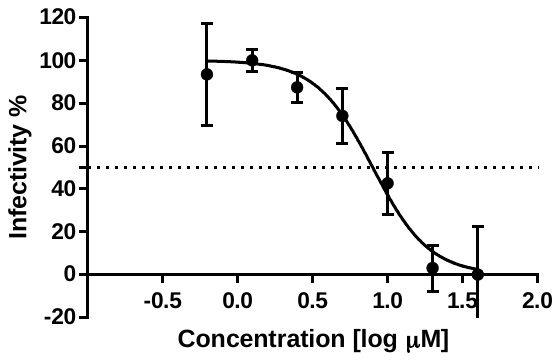

IC_50_=7.97 μM, R^2^=0.93

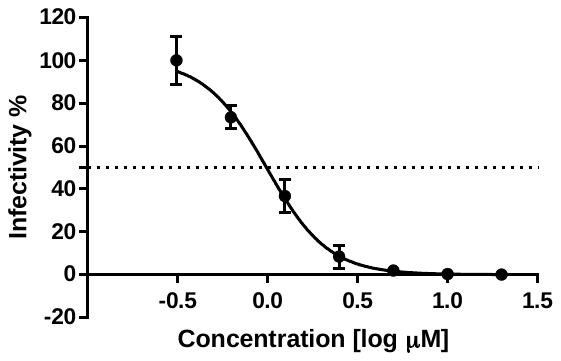

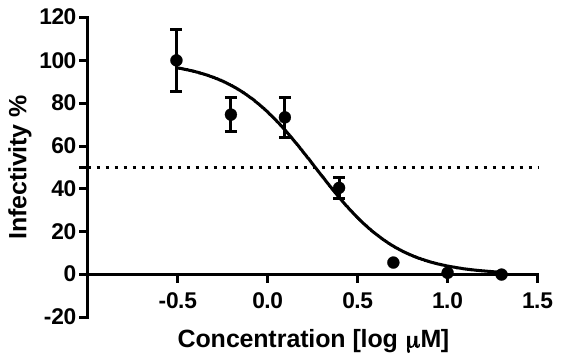

**Delta**

**BA.1**

**BA.2.2**

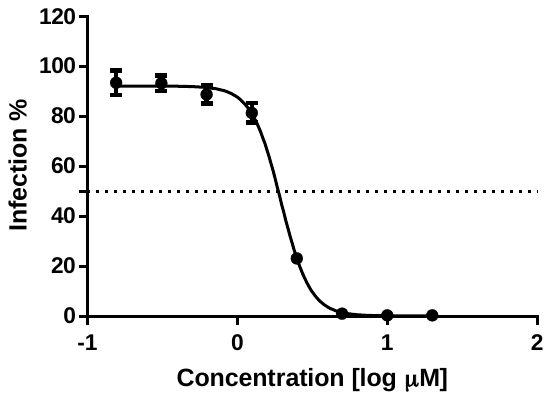

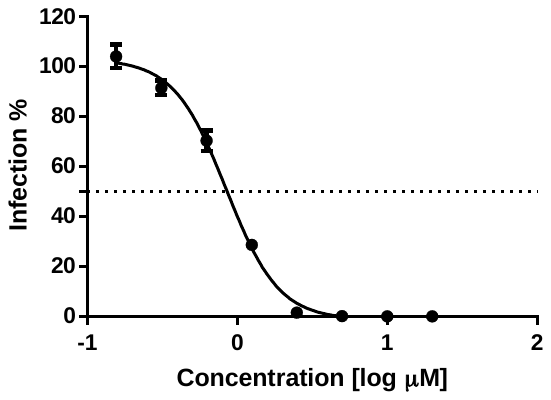

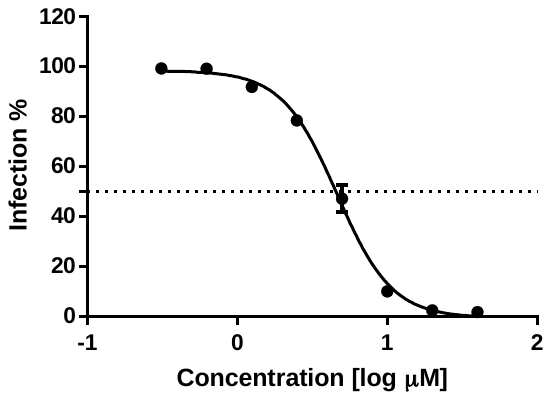

**Remdesivir**

**Molnupiravir**

**Nirmatrelvir**

IC_50_: 1.95 μM, R^2^: 0.99

IC_50_: 0.85 μM, R^2^: 0.99

IC_50_: 4.64 μM, R^2^: 0.99

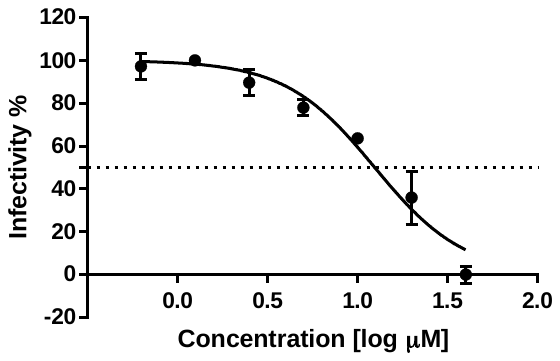

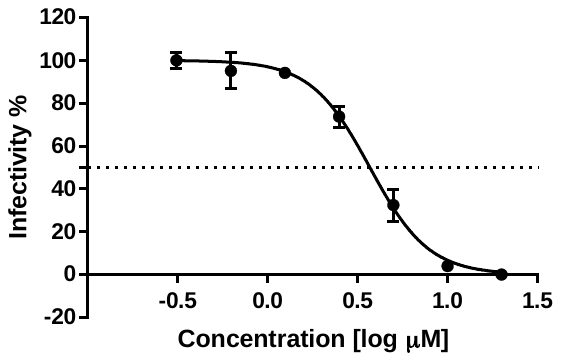

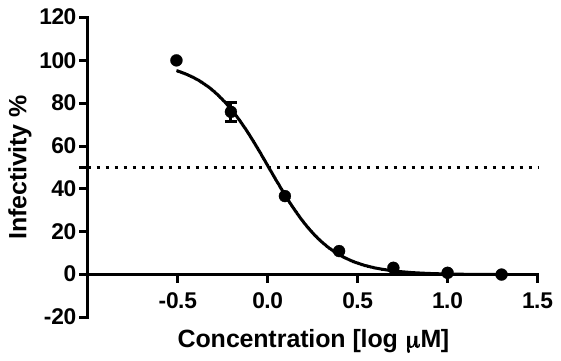

IC_50_=3.70 μM, R^2^=0.98

IC_50_=1.01 μM, R^2^=0.99

IC_50_=12.46 μM, R^2^=0.95

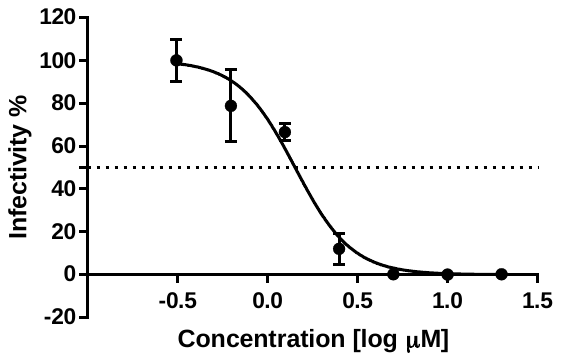

IC_50_: 1.42 μM, R^2^: 0.95

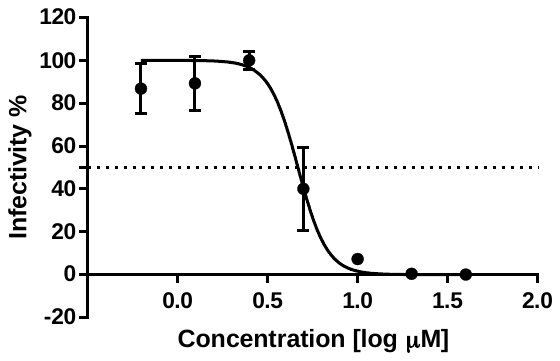

IC_50_: 4.66 μM, R^2^: 0.93

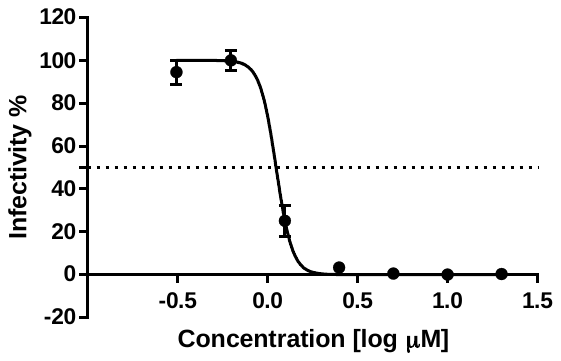

IC_50_: 1.11 μM, R^2^: 0.99

**Remdesivir**

**Molnupiravir**

**Nirmatrelvir**

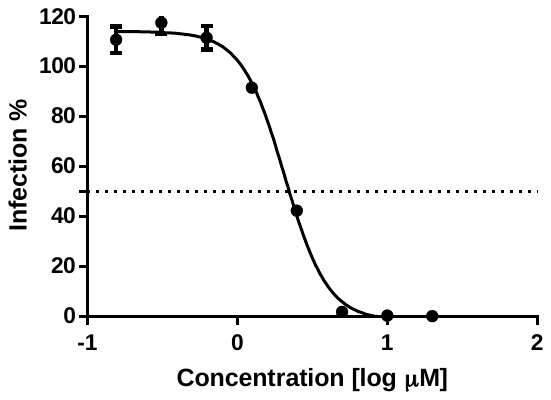

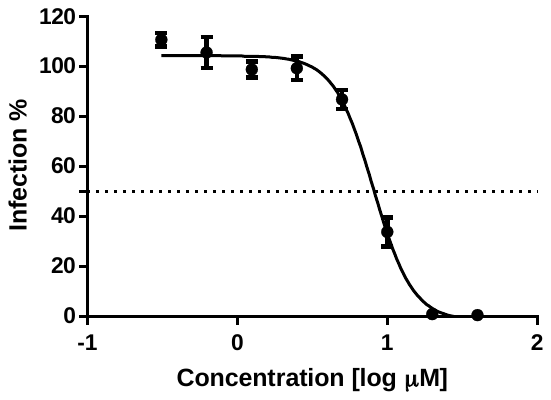

**BA.2**

IC_50_: 2.77 μM, R^2^: 0.96

IC_50_: 2.07 μM, R^2^: 0.99

IC_50_: 8.09 μM, R^2^: 0.98

**BA.2.3**

**BA.2.12.1**

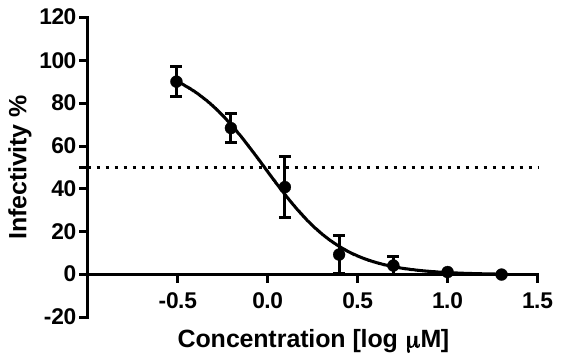

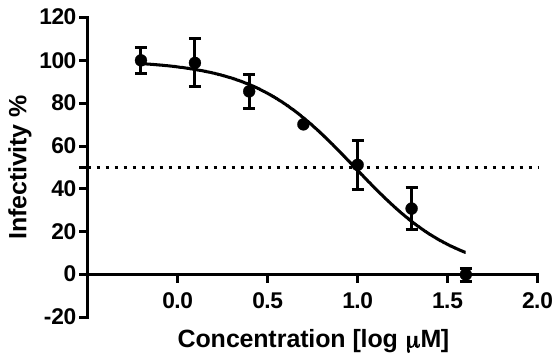

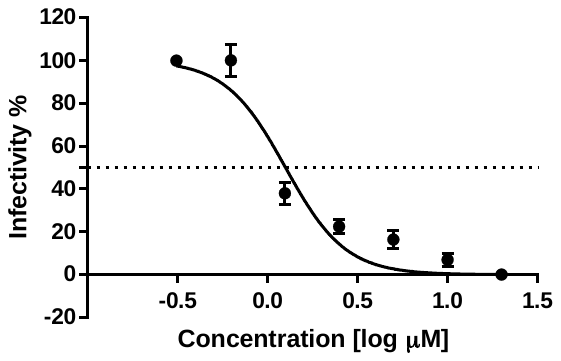

IC_50_: 1.28 μM, R^2^: 0.97

IC_50_: 1.25 μM, R^2^: 0.93

IC_50_: 9.67μM, R^2^: 0.94

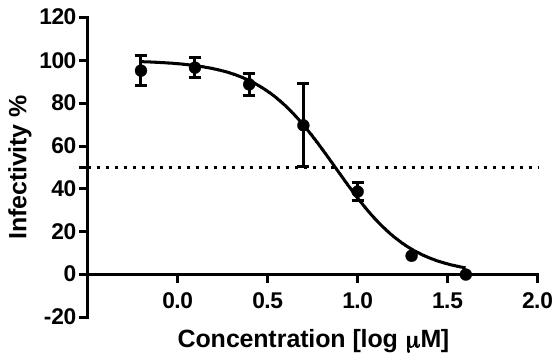

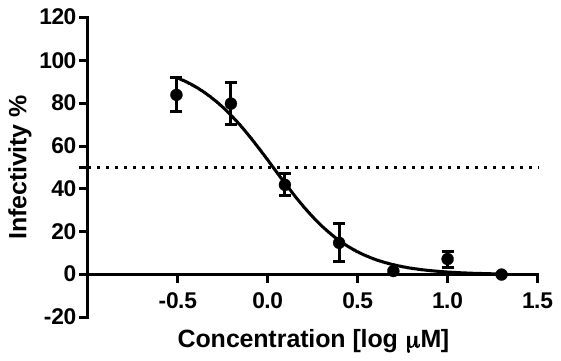

IC_50_: 1.07 μM, R^2^: 0.95

IC_50_: 2.11 μM, R^2^: 0.98

IC_50_: 7.53μM, R^2^: 0.96

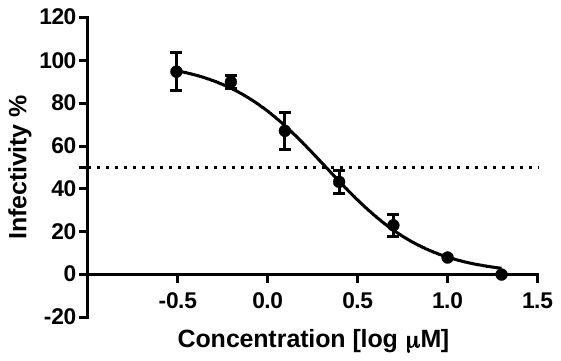

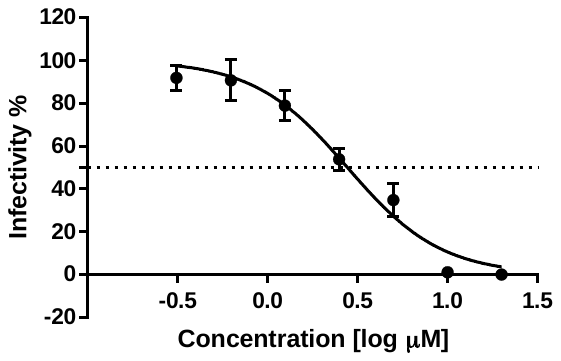

**Remdesivir**

**Molnupiravir**

**Nirmatrelvir**

**BA.4.1.1.**

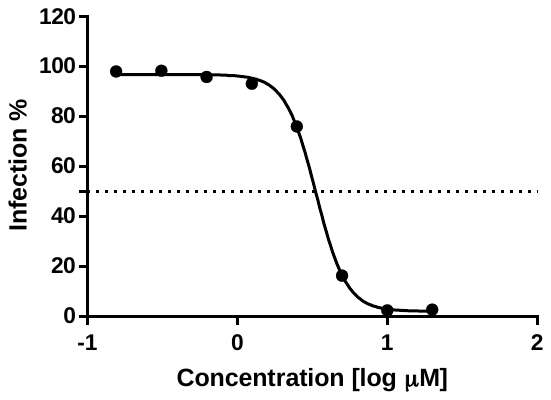

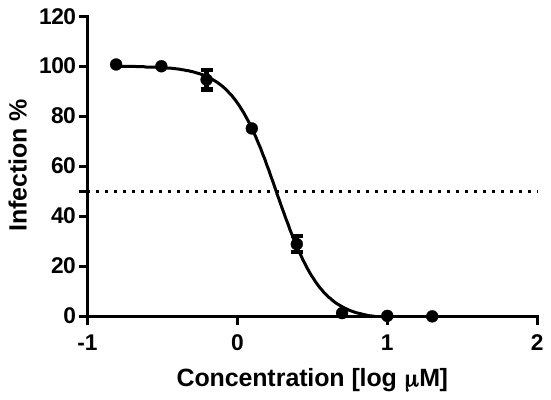

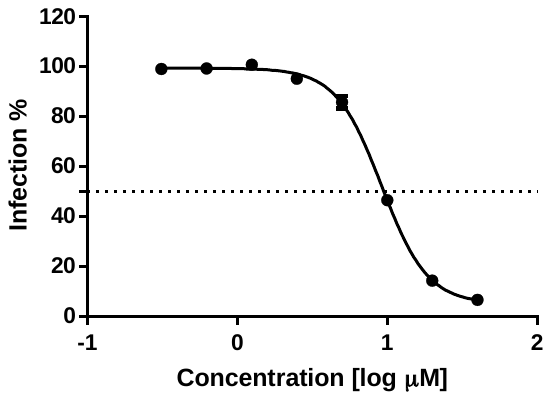

IC_50_: 3.34 μM, R^2^: 0.99

IC_50_: 1.84 μM, R^2^: 0.99

IC_50_: 9.19 μM, R^2^: 0.98

**BA.5.2.1**

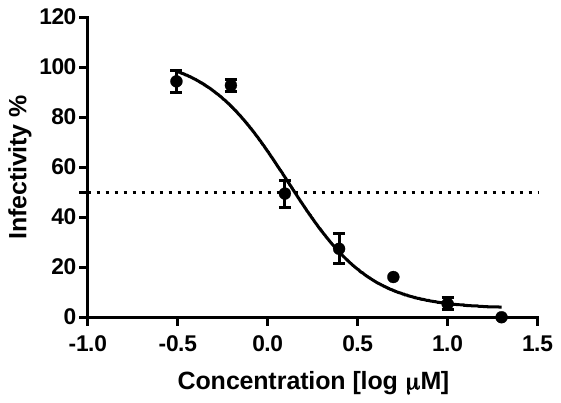

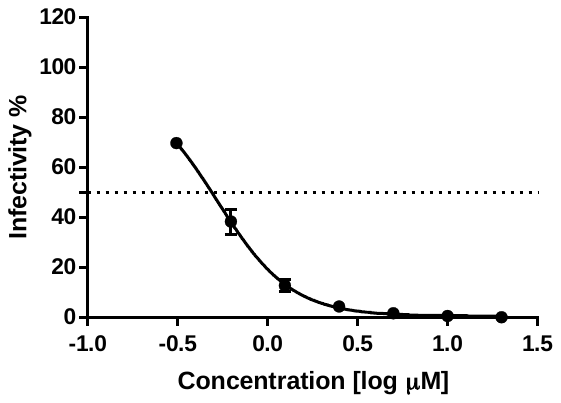

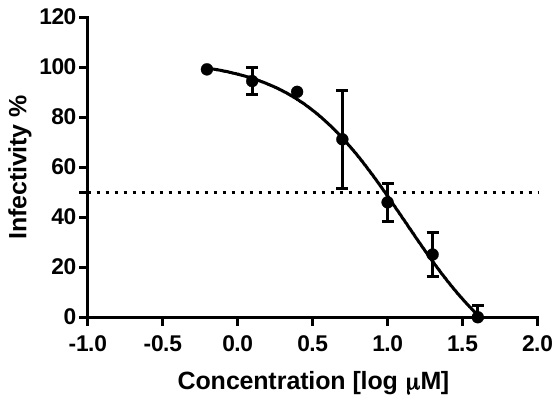

IC_50_=0.54 μM, R^2^=0.99

IC_50_=1.59 μM, R^2^=0.96

IC_50_=7.31 μM, R^2^=0.95

**BA.2.75**

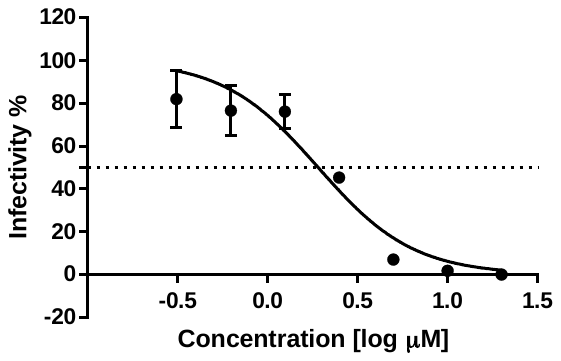

IC_50_=1.91 μM, R^2^=0.91

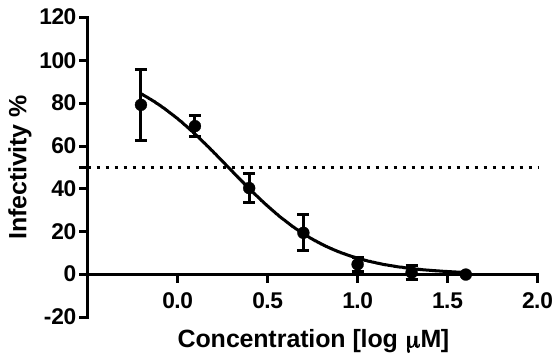

IC_50_=1.92 μM, R^2^=0.94

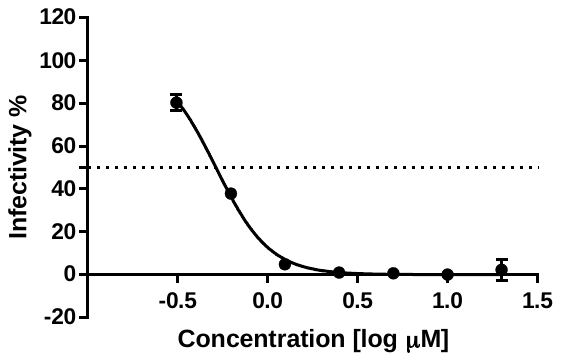

IC_50_=0.51 μM, R^2^=0.99

**Remdesivir**

**Molnupiravir**

**Nirmatrelvir**

**BA.4.6**

**BA.2.75.2**

**BF.7**

IC_50_=4.91 μM, R^2^=0.98

IC_50_=1.15 μM, R^2^=0.99

IC_50_=4.73 μM, R^2^=0.99

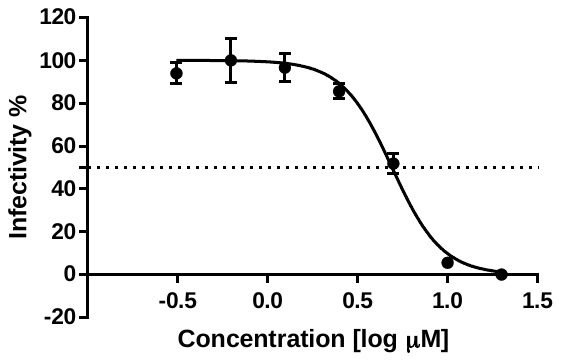

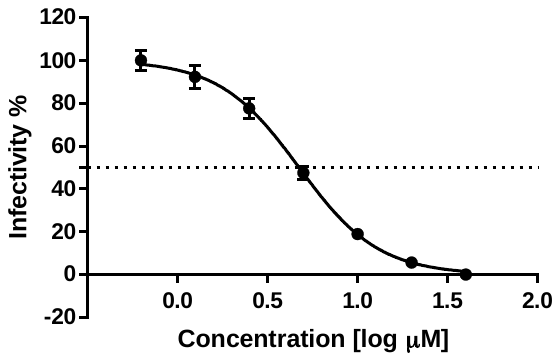

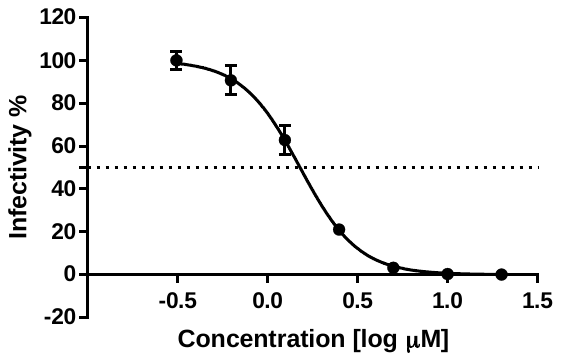

IC_50_=4.34 μM, R^2^=0.98

IC_50_=2.92 μM, R^2^=0.98

IC_50_=1.27 μM, R^2^=0.98

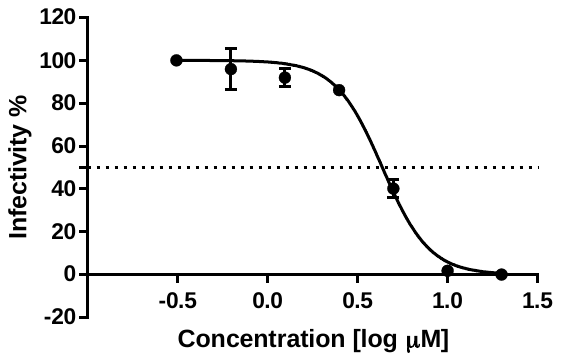

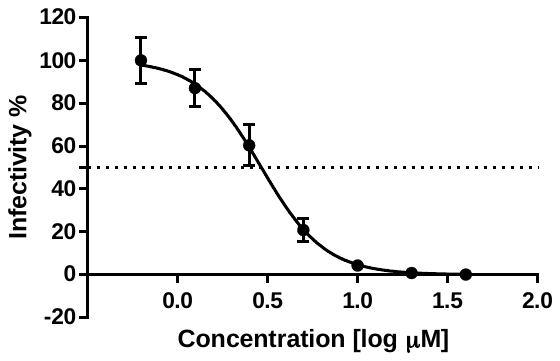

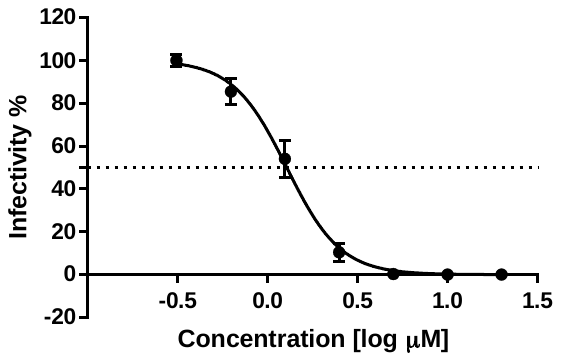

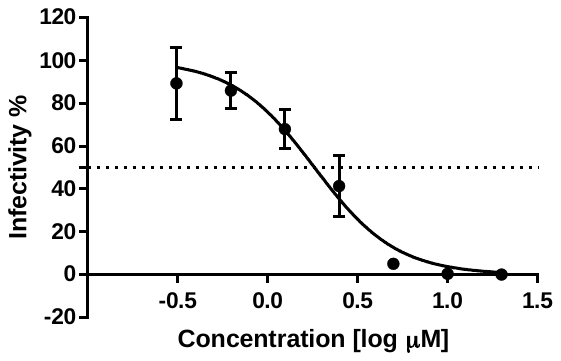

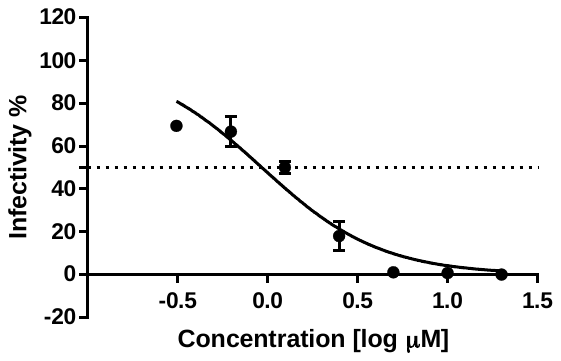

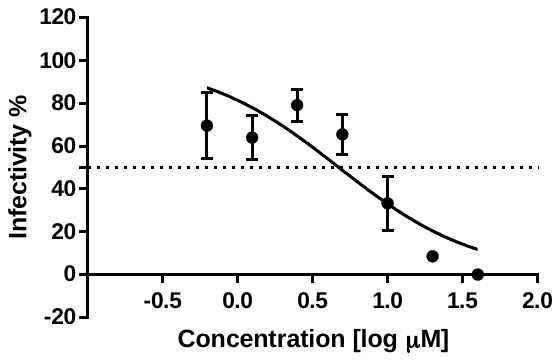

IC_50_=1.82 μM, R^2^=0.94

IC_50_=4.74 μM, R^2^=0.73

IC_50_=0.92 μM, R^2^=0.93

**Remdesivir**

**Molnupiravir**

**Nirmatrelvir**

**BJ.1**

IC_50_=1.32 μM, R^2^=0.96

IC_50_=3.04 μM, R^2^=0.96

IC_50_=4.43 μM, R^2^=0.98

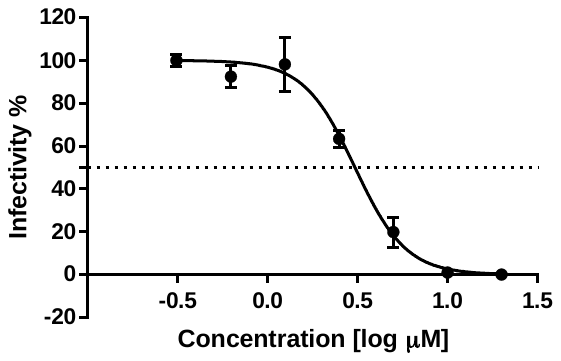

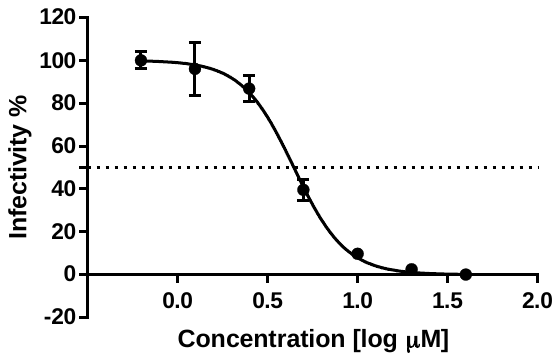

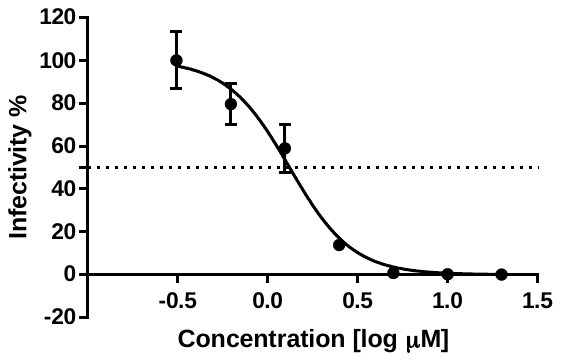

**BQ.1.5**

IC_50_=2.00 μM, R^2^=0.99

IC_50_=6.62 μM, R^2^=0.99

IC_50_=1.30 μM, R^2^=0.97

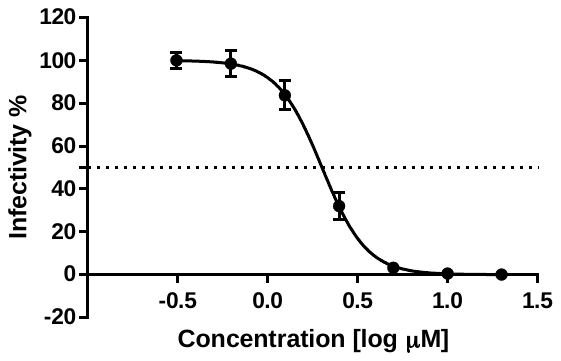

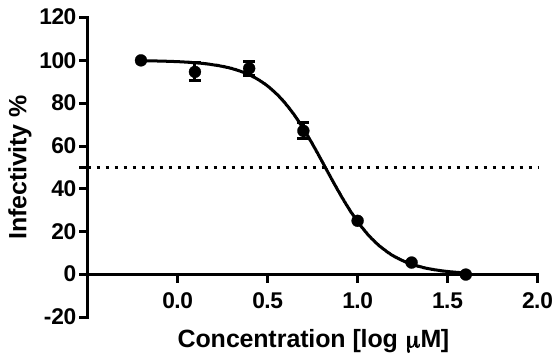

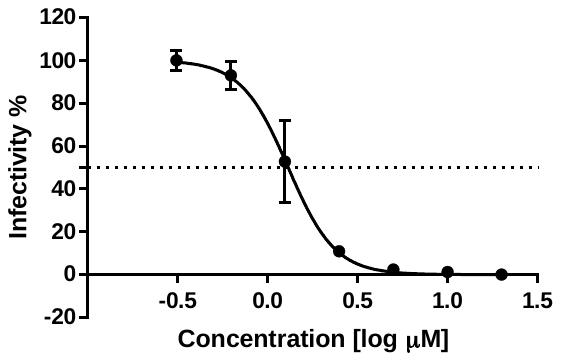

**BQ.1.1**

IC_50_=1.56 μM, R^2^=0.94

IC_50_=1.18 μM, R^2^=0.94

IC_50_=3.22 μM, R^2^=0.82

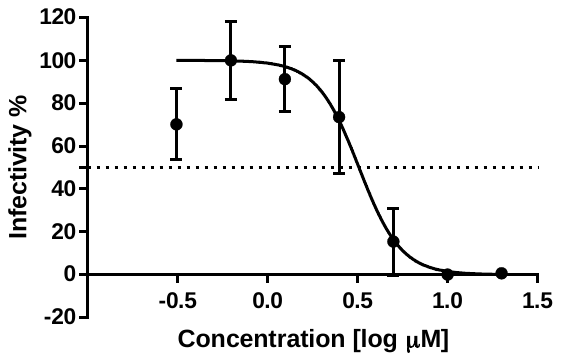

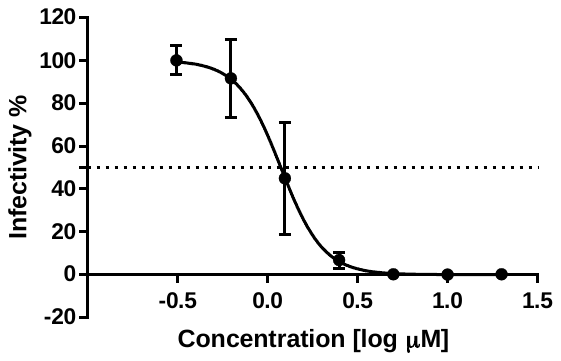

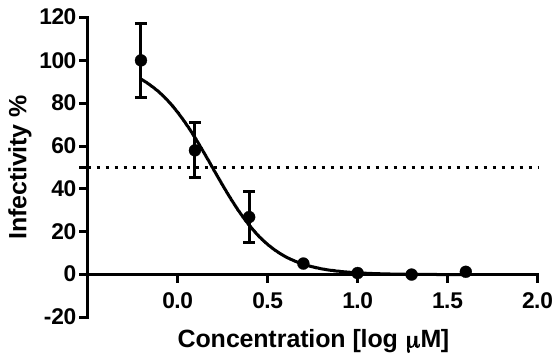

**Remdesivir**

**Molnupiravir**

**Nirmatrelvir**

IC_50_=2.76 μM, R^2^=0.9

IC_50_=3.87 μM, R^2^=0.90

IC_50_=0.94 μM, R^2^=0.93

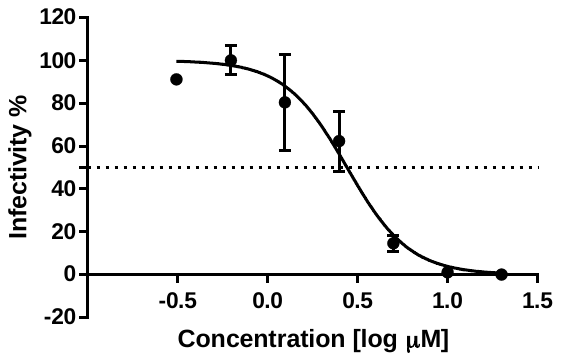

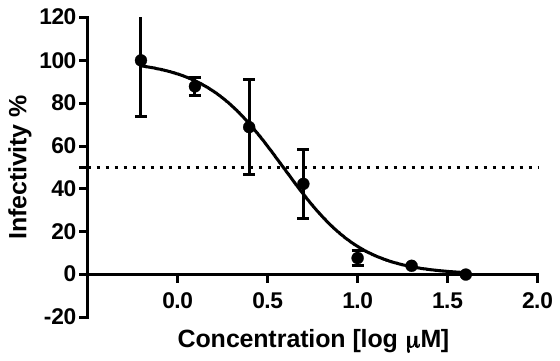

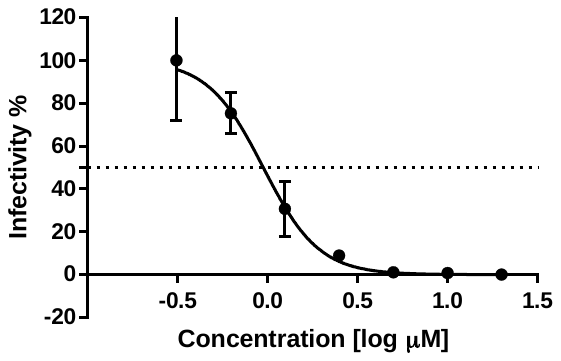

**XBB.1**

**BA.2.75.5**

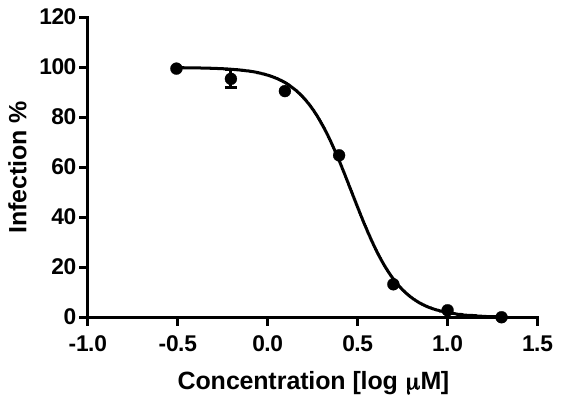

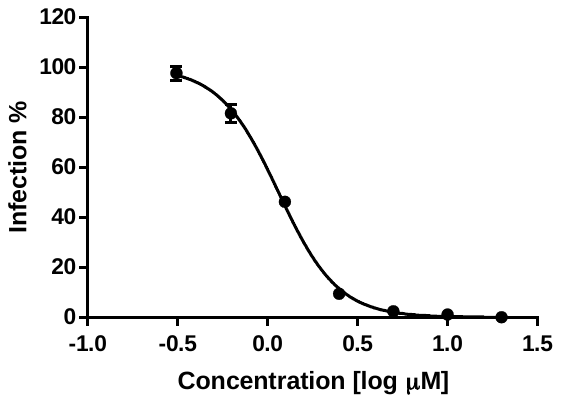

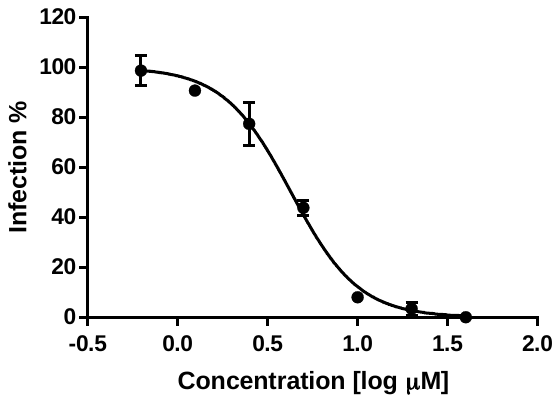

IC_50_=1.15 μM, R^2^=0.96

IC_50_=2.94 μM, R^2^=0.96

IC_50_=4.27 μM, R^2^=0.98

**BA.2.3.20**

IC_50_: 2.15 μM, R^2^: 0.97

IC_50_: 0.83 μM, R^2^: 0.99

IC_50_: 1.92 μM, R^2^: 0.96

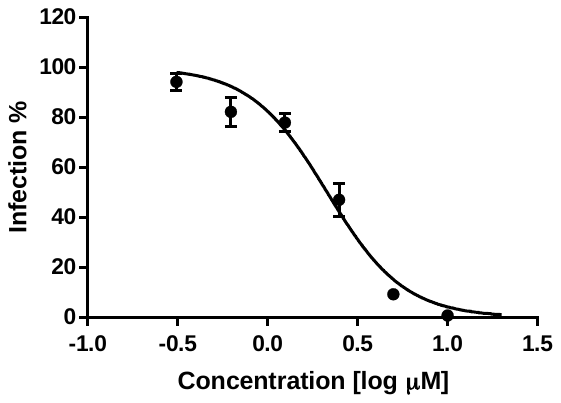

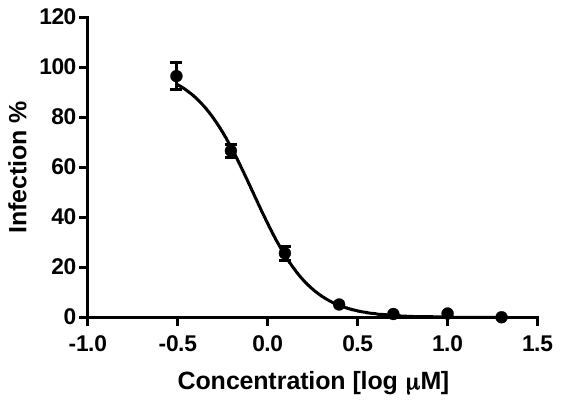

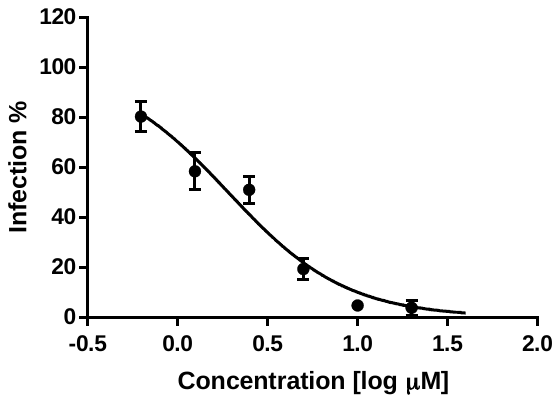

**Remdesivir**

**Molnupiravir**

**Nirmatrelvir**

**XBB**

**XAY.1**

**XBC**

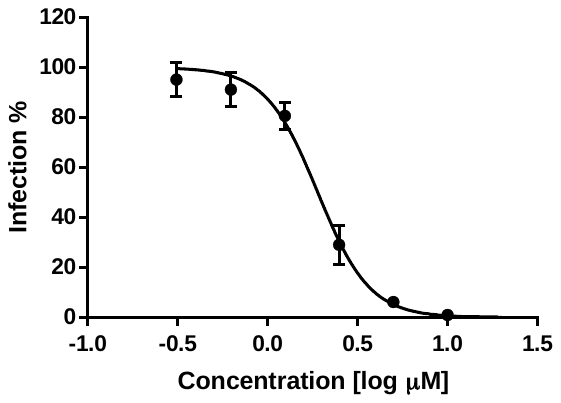

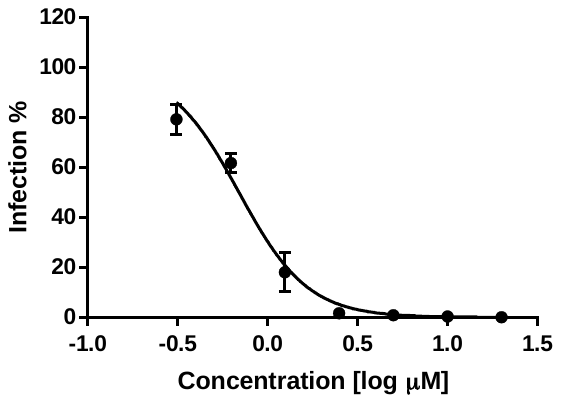

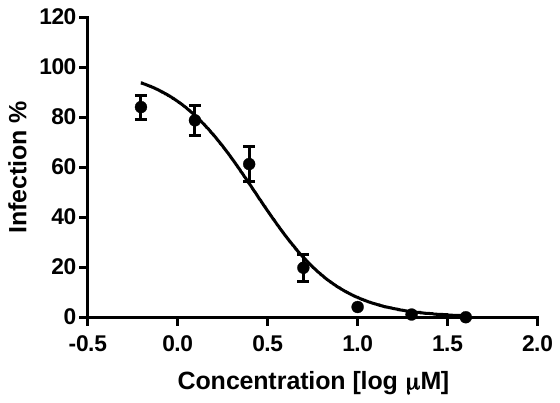

IC_50_: 3.44 μM, R^2^: 0.98

IC_50_: 0.75 μM, R^2^: 0.99

IC_50_: 4.48 μM, R^2^: 0.98

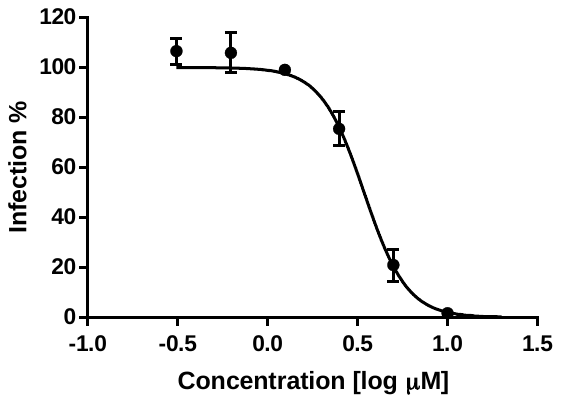

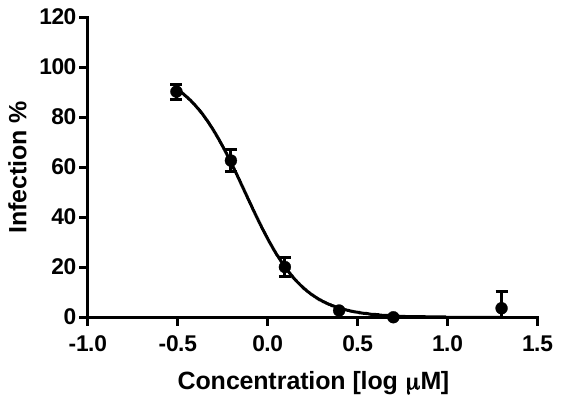

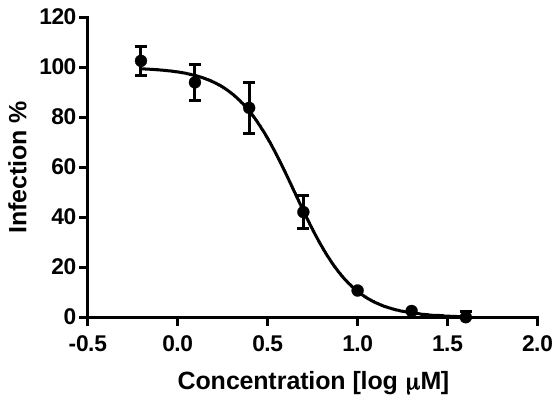

IC_50_: 0.92 μM, R^2^: 0.98

IC_50_: 0.73 μM, R^2^: 0.99

IC_50_: 1.89 μM, R^2^: 0.98

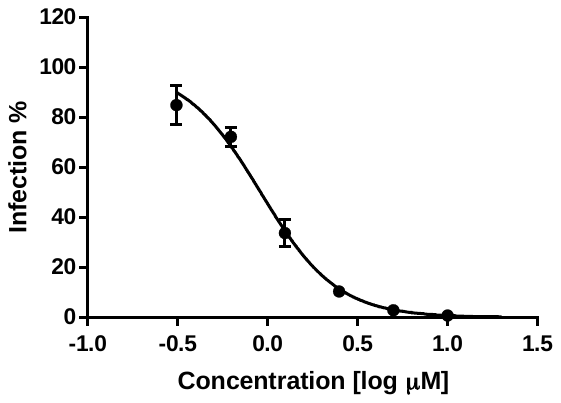

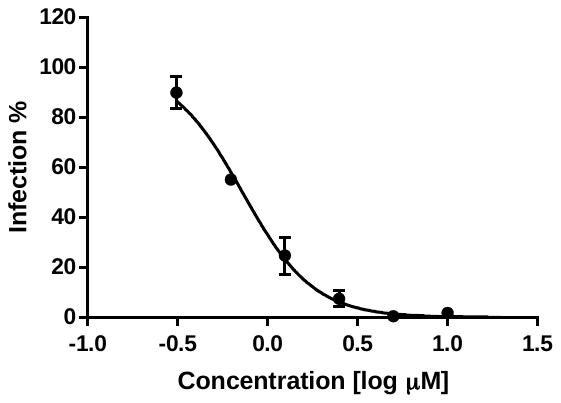

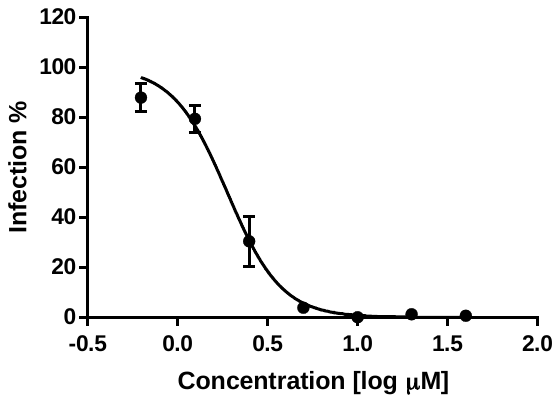

IC_50_: 1.90 μM, R^2^: 0.98

IC_50_: 0.69 μM, R^2^: 0.97

IC_50_: 2.70 μM, R^2^: 0.96

**Remdesivir**

**Molnupiravir**

**Nirmatrelvir**

**BN.1.9**

IC_50_: 2.19 μM, R^2^: 0.94

IC_50_: 0.59 μM, R^2^: 0.98

IC_50_: 2.36 μM, R^2^: 0.97

**BN.1.5**

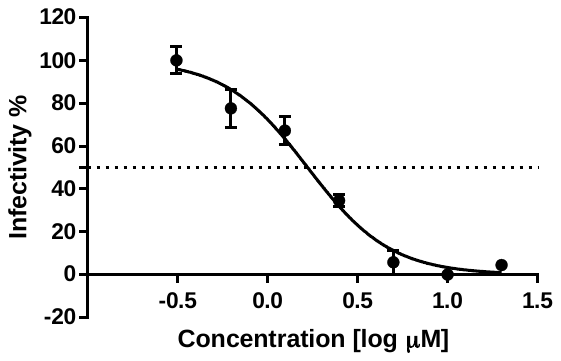

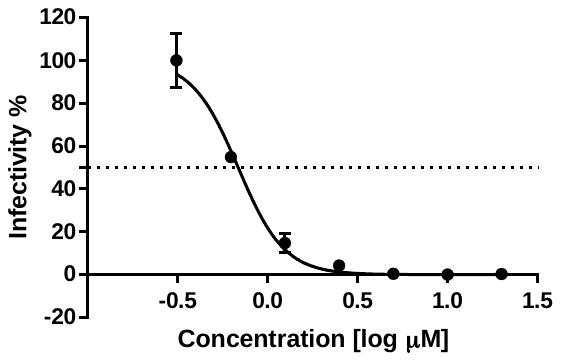

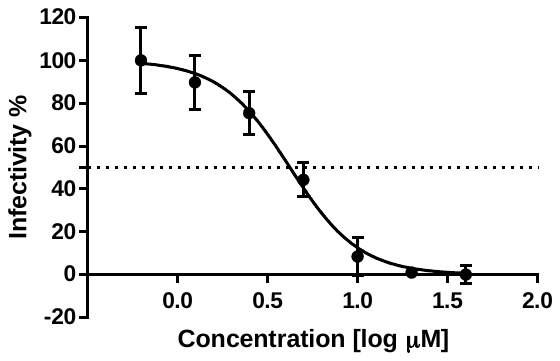

IC_50_: 1.67 μM, R^2^: 0.96

IC_50_: 0.69 μM, R^2^: 0.97

IC_50_: 4.21 μM, R^2^: 0.95

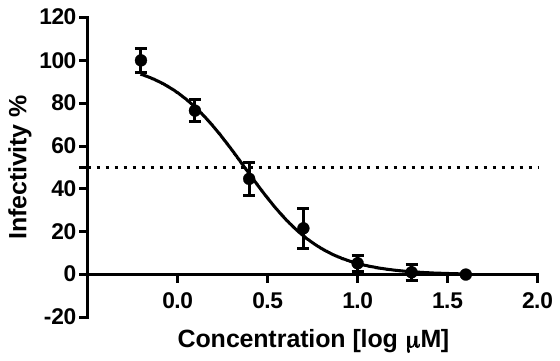

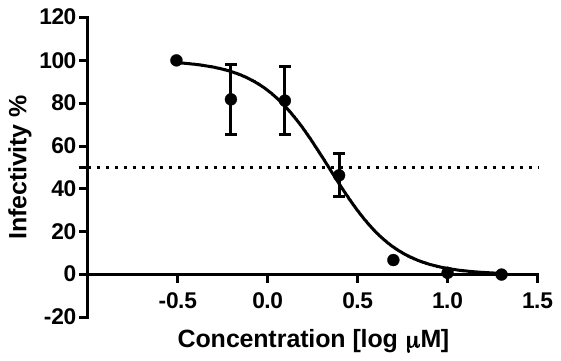

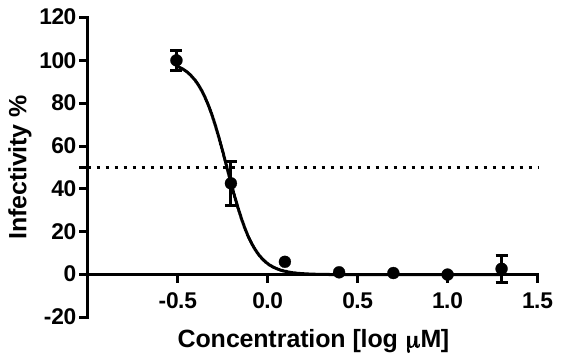

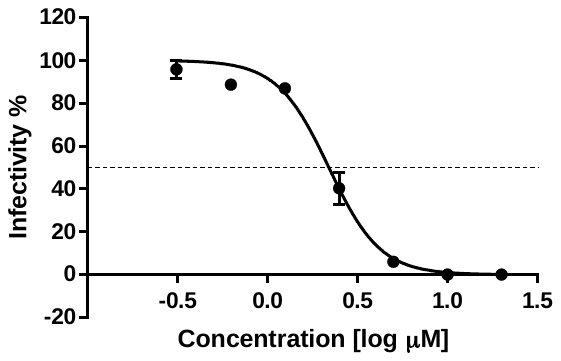

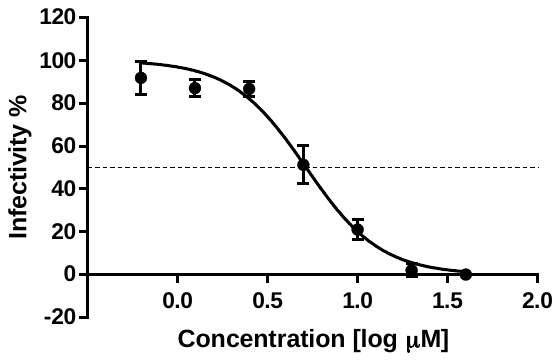

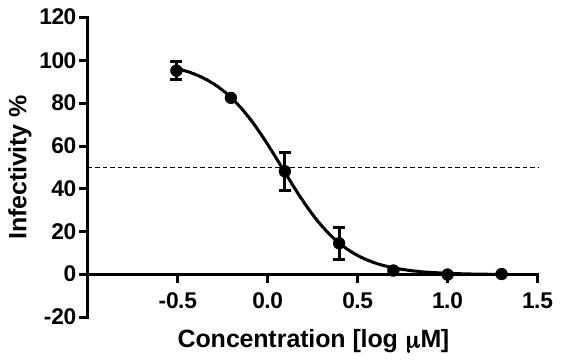

IC_50_: 2.19 μM, R^2^: 0.98

IC_50_: 1.20 μM, R^2^: 0.98

IC_50_: 5.18 μM, R^2^: 0.97

**XBB.1.5**

Supplementary Table S1

Supplemental Table 1. List of SARS-CoV-2 variants used in this study.

| Variant (Lineage) | GISAID  ID | Substitution  Nsp 5 Nsp 12 | | Virus name |
| --- | --- | --- | --- | --- |
| Reference (A) | EPI_ISL_407193 |  | | hCoV-19/South Korea/KCDC03/2020 |
| Alpha (B.1.1.7) | EPI_ISL_738139 | - | P323L | hCoV-19/South Korea/KDCA0001/2020 |
| Beta (B.1.351) | EPI_ISL_762992 | K90R | P323L | hCoV-19/South Korea/KDCA0463/2020 |
| Gamma (P.1) | EPI_ISL_1622497 | - | P323L | hCoV-19/South Korea/KDCA2945/2021 |
| Delta (B.1.617.2) | EPI_ISL_2887353 | - | P323L, G671S | hCoV-19/South Korea/KDCA5439/2021 |
| BA.1 (B.1.1.529) | EPI_ISL_6959993 | P132H | P323L | hCoV-19/South Korea/KDCA18126/2021 |
| BA.2 | EPI_ISL_13086512 | P132H | P323L | hCoV-19/South Korea/KDCA61368/2022 |
| BA.2.2 | EPI_ISL_8885887 | P132H | P323L | hCoV-19/South Korea/KDCA26119/2021 |
| BA.2.12.1 | EPI_ISL_13086514 | P132H | P323L | hCoV-19/South Korea/KDCA58217/2022 |
| BA.2.3 | EPI_ISL_13086513 | P132H | P323L | hCoV-19/South Korea/KDCA61369/2022 |
| BA.4.1.1 | EPI_ISL_13086515 | P132H | P323L | hCoV-19/South Korea/KDCA61370/2022 |
| BA.5.2.1 | EPI_ISL_13086516 | P132H | P323L | hCoV-19/South Korea/KDCA61371/2022 |
| BA.2.75 | EPI_ISL 14507613 | P132H | P323L, V405F, G671S | hCoV-19/South Korea/KDCA96765/2022 |
| BA.4.6 | EPI_ISL_14780352 | P132H | P323L | hCoV-19/South Korea/KDCA103718/2022 |
| BA.2.75.2 | EPI_ISL_15315727 | P132H | P323L, G671S | hCoV-19/South Korea/KDCA130199/2022 |
| BF.7 | EPI_ISL_15535610 | P132H | P323L | hCoV-19/South Korea/KDCA137894/2022 |
| BQ.1.5 | EPI ISL 15535612 | P132H | P323L, Y273H | hCoV-19/South Korea/KDCA137896/2022 |
| BQ.1.1 | EPI ISL 15848736 | P132H | P323L, Y273H | hCoV-19/South Korea/KDCA155725/2022 |
| BJ.1 | EPI ISL 15535611 | P132H | P323L, G671S | hCoV-19/South Korea/KDCA137895/2022 |
| XBB.1 | EPI_ISL_15695954 | P132H | P323L, G671S | hCoV-19/South Korea/KDCA152302/2022 |
| BA.2.3.20 | EPI_ISL_16299727 | P132H | P323L | hCoV-19/South Korea/KDCA166997/2022 |
| BA.2.75.5 | EPI_ISL_16299728 | P132H | P323L, G671S | hCoV-19/South Korea/KDCA166998/2022 |
| XBB | EPI_ISL_16299731 | K90R, P132H | P323L, G671S | hCoV-19/South Korea/KDCA167001/2022 |
| XAY.1 | EPI_ISL_16299730 | P132H | P323L, A529V, G671S | hCoV-19/South Korea/KDCA167000/2022 |
| XBC | EPI_ISL_15315728 | - | P323L, A529V, G671S, ins823D | hCoV-19/South Korea/KDCA130200/2022 |
| BN.1.5 | EPI_ISL_16852231 | P132H | P323L, G671S | hCoV-19/South Korea/KDCA0001/2023 |
| XBB.1.5 | EPI_ISL_16852232 | P132H | P323L, G671S | hCoV-19/South Korea/KDCA0002/2023 |
| BN.1.9 | EPI_ISL_17171894 | P132H | P323L, G671S | hCoV-19/South Korea/KDCA218115/2023 |
